# A uniform tissue-clearing framework and mesoSPIM-ultra enable cm-scale single-neuron tracing

**DOI:** 10.64898/2026.06.29.734841

**Authors:** Marko Pende, Jared M. Cregg, Saiedeh Saghafi, Samuel Broadbent, Alma Avdibasic, Johannes Roeles, Sofia-Christina Papadopoulos, Ryan P. Seaman, Nika Pende, Marta Solano Mateos, Katelyn Jamwal, Melody Wunch, Pawel Pasierbek, Alberto Moreno-Cencerrado, Solomiia Korchynska, Romana Hauer, Paul Anderson, Paul Supper, Maria Eleni Kastriti, Daniel Reumann, Michael Moorhead, Joel H. Graber, Petra Scholze, Julia U. Henschke, Eike Budinger, Jürgen A. Knoblich, Thomas Klausberger, Igor Adameyko, Tibor Harkany, Vivek Kumar, Mary Teena Joy, Ole Kiehn, Hans-Ulrich Dodt, Fabian F. Voigt, Prayag Murawala

**Author notes:** These authors contributed equally (Marko Pende, Jared M. Cregg).

## Abstract

Tissue-clearing and light-sheet microscopy have transformed volumetric imaging of intact organs, yet limited mechanistic understanding of dehydration-based clearing continues to constrain rational protocol design and broader applicability. Here, we define the cardinal chemical and physical principles underlying dehydration-based tissue-clearing and establish a new pipeline for large-volume imaging. To maximize imaging performance, we developed the “mesoSPIM-ultra”, an upgraded mesoSPIM platform with a temperature-controlled sample chamber, a large field-of-view (FoV) camera and specialized optics to achieve long-working-distance, high-resolution imaging of cleared samples. We applied this approach to investigate the projectome of *Chx10*^+^ neurons, a cell population with complex axonal morphologies along the entire mouse spinal-cord and brain, and implicated in ipsilateral orienting behaviors. By combining behavioral analysis with *post-hoc* single-neuron reconstructions, we revealed previously inaccessible branching architectures and long-range projections extending from the brainstem to the spinal cord. Together, our work establishes a mechanistic foundation for tissue-clearing and scalable imaging.

## Introduction

Understanding how neural circuits generate locomotion and coordinate behavior requires mapping of neuronal connectivity across the entire nervous system. Despite major advances in projectomics, reconstructing long-range axons that course along the brain and spinal cord remains a significant challenge. As a result, fundamental questions - such as how distributed circuits coordinate movement, how descending pathways interact with spinal networks, or how circuit architecture gives rise to behavior - remain incompletely understood. Traditional histological sectioning provides detailed structural information but is labor-intensive and prone to section loss and deformation, thereby limiting reconstruction of large-scale neuronal networks. Consequently, obtaining comprehensive maps of neural circuits across the intact central nervous system remains difficult. In contrast, single-photon, two-photon, and light-sheet fluorescence microscopy (LSFM) enable non-destructive three dimensional (3D) optical sectioning but are limited by light scattering in intact tissues.

Tissue-clearing methods can overcome these challenges by rendering samples optically transparent^1–3^. Numerous clearing protocols have been developed, including aqueous-based^4–18^ and dehydration-based organic-solvent-dependent^19–28^ protocols. While aqueous methods preserve endogenous protein-based fluorescence, dehydration-based methods offer faster clearing and higher transparency, particularly for dense or lipid-rich tissues. Protocols that shrink the sample reduce the imaging volume^24–26,28,29^ and facilitate whole-organ visualization, whereas methods that preserve or expand sample improve the resolution^17,18,30,31^. Moreover, tissue composition itself strongly influences clearing outcomes, e.g., differences in lipid content between young and aged tissue, or between organs can impact the transparency, morphology, and autofluorescence. Despite the large number of clearing protocols available, the choice of the appropriate protocol for a given tissue is hindered by the lack of mechanistic understanding and the standardized evaluation of clearing performance. Further, while advances in imaging, tissue-clearing and sample analysis have offered insights into 3D tissue organization^32^, developmental processes^3^, organ pathology^33^, neural connectivity^1^, and regenerative mechanisms^9,34^, no tissue-clearing workflows have been published to date for imaging multi-cm long-range axonal projections at the spatial scale necessary to link, e.g., hindbrain neurons that project in the spinal cord. This is surprising given the central role of long-range connectivity in locomotion, where alternating limb movements are coordinated by neural projections extending centimeters in mice and up to meters in humans.

To address these limitations, we assessed existing clearing methods; examined the chemical and physical parameters underlying dehydration-based tissue-clearing and developed optimized clearing strategies that preserve fluorescence and structural integrity in mineralized and lipid-rich tissues. Furthermore, we developed the mesoSPIM-ultra light-sheet microscope featuring improved optics, a larger FoV camera and a temperature-controlled sample chamber for optimized transparency enabling long-range neuronal tracing from the brain to the sacral of the spinal cord. Finally, by combining a physicochemically informed tissue-clearing approach with the mesoSPIM-ultra-based imaging, we resolved the complex arborization of axonal projections of *Chx10^+^*movement-orienting neurons in the intact mouse central nervous system (CNS), revealing hitherto unrecognized anatomical heterogeneity that offers unique projectome relationships in motor circuits. Together, these advances establish a reproducible framework for high-fidelity whole-scale mapping of the structure and connectivity of biological components at mesoscale biological systems.

## RESULTS

### Quantitative Analysis of Tissue Transparency

We sought to develop a quantitative approach to assess tissue transparency and compare the clearing performance of different protocols. As described by Lambert’s law, transparency of a biological sample is determined by total transmission (TT), the scattering of light (haze; H), and the optical properties of the medium (sample) through which light propagates, (T = I/I_0_ = e^-kx^, whereas H = I_d_/I; where T=transmission, I=transmitted light intensity, I_d_=scattered light intensity, I_0_=incident light intensity, k=absorption co-efficient, and x=thickness of the material). Despite this quantitative basis for defining transparency, most tissue-clearing methods rely on qualitative evaluation, via visual inspection on graph paper or by using a US Air Force (USAF) resolution chart to demonstrate optical transparency. However, visualization of a cleared tissue on the USAF chart provides only an approximate indication of TT, with no information on H. In addition, differences in the placement of a sample with respect to the USAF chart can impact visual assessment (Figure 1A). Thus, an informed assessment of transparency must consider all parameters described by Lambert’s law as well as H, and the refractive index (RI) properties of the sample and its surrounding medium.

**Figure 1.**
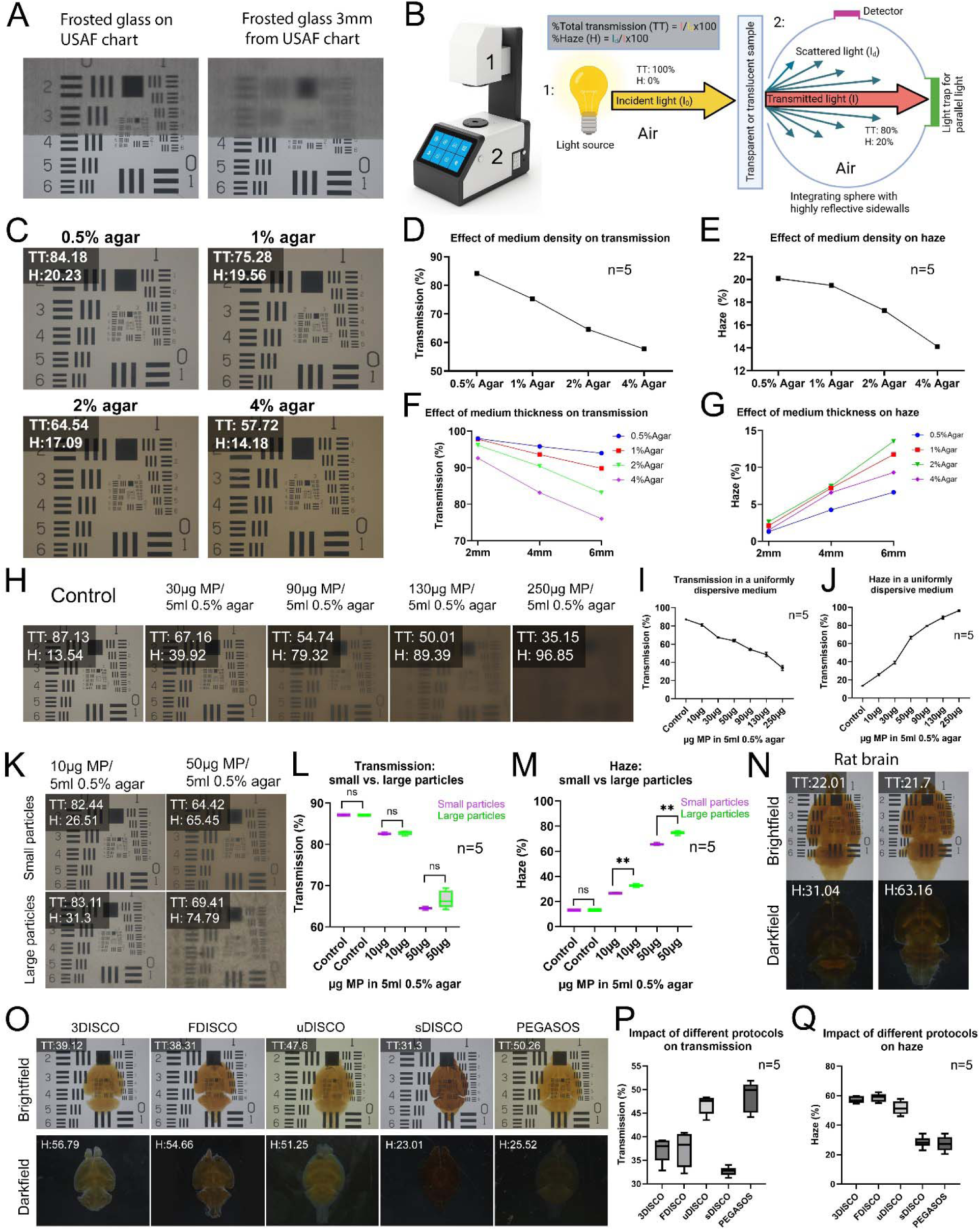
Quantifying Transparency through Haze and Transmission. (A) Increased optical path length amplifies light scattering and its consequent effect on transparency, demonstrated using a frosted slide and USAF chart. (B) Schematic illustrating light propagation through air followed by a transparent or translucent sample. Transparency was quantified by measuring total transmission (TT) and haze (H) with a hazemeter. (C-E) The impact of medium density, i.e., agar concentration, on TT and H. Results shown in C are plotted in (D, E) (n = 5 times measured). (F, G) The combined impact of medium density (agar concentration) and medium thickness on (F) TT and (G) H. (H-J) Impact of the density of scattering particles, i.e., milk powder (MP), on TT and H. Results shown in H are plotted in (I, J) (n = 5 times measured).(K-M) Impact of light scattering on TT and H, using different particle sizes and concentrations to simulate differences in light scattering (L, M: ns: not significant, P > 0.05, ** P ≤ 0.01, Mann-Whitney test). Results shown in K are plotted in (L, M) (n = 5 times measured). (N) Rat brains with different extents of lipid extraction illustrate the contribution of tissue scattering to TT and H. Brains were incubated in THF for 1-week (right sample) or 2-weeks (left sample) before assessment. (O-Q) Impact of different dehydration-based tissue-clearing protocols on TT and H, using mouse brains as samples. Boxplots (P, O) show results from O plotted (n = 5 brains).

To move beyond conventional qualitative assessment, we simultaneously quantified TT and H in tissue-cleared samples by employing a commercial hazemeter, a device routinely used to measure the transparency of glass or plastic in relevant industries (Figure 1B). Our aim was to determine how physical parameters, sample density, thickness, and particulate content (e.g., residual lipids) influence the TT and H. We first validated the hazemeter using agar gels of four different densities (Figure 1C). As expected, TT decreased steadily with increasing agar density (Figure 1D), consistent with greater optical attenuation. Further, H also decreased with increasing density (Figure 1E), suggesting that RI homogeneity in denser gels reduces scattering sites. We then evaluated the effect of gel (medium) thickness on TT. As expected, we observed that increased thickness was associated with decreased TT (Figure 1F), and increased H at all four agar concentrations (Figure 1G). The latter finding indicates that thicker media provides more opportunities for light scattering.

Light scattering fundamentally shapes light-tissue interactions. Effective tissue-clearing requires identifying and addressing the sources of scattering including residual lipids and RI mismatches, which limit transparency and imaging depth. To model scattering caused by uniformly dispersed particles, we first dissolved milk powder (MP) at various concentrations in water and then created 0.5% low-melting agar gels of identical thickness (4 mm) (Figure 1H). Increasing MP concentration caused a progressive decrease in TT and a corresponding increase in H (Figures 1I-J). To simulate scattering local variations in particle density, we prepared 0.5% agar gels in which MP was only coarsely mixed, creating non-uniform clusters of larger particles, and compared these to uniformly distributed smaller particles generated by dissolving MP of the same amount of agar. Compared to gels with uniformly distributed smaller particles, gels containing coarsely mixed MP showed significantly increased H, but TT remained unchanged (Figures 1K-M). Collectively, these findings demonstrate that TT and H are not linearly related; rather, they depend on the sample’s microstructural and RI heterogeneity—parameters directly affected by the tissue-clearing chemistry and immersion medium.

Although brightfield imaging of cleared samples via USAF chart remains the standard qualitative method to demonstrate tissue transparency, it provides only a qualitative TT value, and the readout is constrained by the smallest resolvable element on the chart, making it impossible to distinguish differences in overall sample transparency (Figure 1C). To illustrate these limitations, we compared two rat brains that had been dehydrated for different lengths of time in tetrahydrofuran (THF) and were then RI-matched with dibenzyl ether (DBE), resulting in samples differing in residual lipid content. The cleared rat brains were fully immersed, together with the USAF chart, in RI medium and imaged under brightfield conditions. Despite their different clearing histories, both samples produced identical maximum resolution values on the USAF chart and similar TT values using the hazemeter (Figure 1N). However, the H values of the two samples measured via hazemeter exhibited distinct scattering properties (Figure 1N). Darkfield microscopy, which relies on scattered light for image formation, offers a natural qualitative proxy for haze, allowing for a more comprehensive assessment of clearing efficiency in comparison to a USAF chart when a hazemeter is not available. We then compared the impact of five common dehydration-based clearing protocols^21,23,24,26,28^, on optical properties of (i.e., TT and H) tissue-cleared mouse brains (Figures 1O-Q). The strong positive correlation between H values and darkfield intensity, and between TT values and brightfield transmission, supports the complementary nature of these parameters in evaluating tissue transparency.

Altogether, our results demonstrate that quantitative assessment of tissue transparency must include both TT and H values, as they provide distinct yet interrelated insights into how light interacts with biological samples and clearing media. It is noteworthy that these values can also be influenced by the intrinsic properties of the sample itself, e.g., tissue type, cellular density, and lipid content, such that TT and H comparisons are most meaningful within the same sample category, as structurally distinct tissues such as kidney *vs.* brain could yield different values even under identical clearing conditions.

### Comparison of Established Clearing Protocols and Evaluation of Dehydration Agents for Autofluorescence Quenching and Fluorescent Protein Preservation

To better define the parameters governing tissue-clearing and fluorescent protein (XFP) stability, we systematically benchmarked five widely used organic solvent-based clearing protocols, i.e., 3DISCO^21^, FDISCO^24^, sDISCO^26^, uDISCO^23^, and PEGASOS^28^, for their ability to preserve fluorescence from Thy1-YFP-H and CAGGs:tdTomato mouse-brain tissue across each step of the workflow (Figure S1A-T). Notably, we implemented one modification to the previously published sDISCO protocol^23^, which uses propyl gallate (PG) to stabilize DBE (sDBE). Thereby sDISCO intends to prevent peroxide-mediated XFP quenching (Figure S1G) and improves long-term fluorophore stability (Figure S1H). However, sDBE can lead to precipitate formation, likely due to interactions of PG with heme groups from residual blood remaining after incomplete perfusion (Figures S1I,J). To minimize precipitation, we replaced THF with tert-butanol (tB) during the early steps of dehydration. This modification did not affect XFP signals, even at 90% concentration (Figure S1K), underscoring a role for tB in preventing precipitate formation.

Next, we quantified the change of the mean and maximum fluorescence intensity relative to the initial signal measured after fixation in 4% PFA and PBS washing (Figures S1B,C,E,F,M,N,P,Q,S,T). Across all protocols, fluorescence intensity progressively declined during dehydration (Figure S1). Among the five protocols tested, 3DISCO, FDISCO, and sDISCO use THF as the primary dehydration agent, whereas uDISCO and PEGASOS rely on tB. Overall, THF-based protocols (Figures S1A,D,G) caused greater fluorescence loss than tB-based approaches (Figures S1O,R). For GFP, signal reduction was highest in 3DISCO and sDISCO, whereas FDISCO showed improved preservation despite the presence of THF. uDISCO and PEGASOS exhibited lower GFP loss overall (Figure S1). TdTomato fluorescence was substantially more sensitive, particularly in sDISCO, while PEGASOS and uDISCO again showed the strongest preservation (Figure S1). Interestingly, in most protocols, fluorescence transiently decreased during intermediate dehydration steps (50–80%) but partially recovered at 90% solvent concentration. Among all protocols, PEGASOS achieved the best overall fluorescence preservation after dehydration (Figures S1R–T), likely due to PEG-containing final dehydration steps that retain residual water and stabilize endogenous fluorophores.

We also assessed protocol-autofluorescence occurrence, as it directly impacts the signal-to-noise ratio (SNR) during imaging. Overall, THF-based protocols slightly reduced tissue autofluorescence, whereas tB-based protocols produced a modest increase; however, in all cases these changes remained below 0.5-fold in both mean and maximum intensity measurements at 488 nm and 568 nm excitation wavelengths (Figure S1). The use of DBE-based RI-matching media further reduced autofluorescence in both THF- and tB-based clearing strategies. In contrast, the RI media used in uDISCO and PEGASOS, namely 1:2 benzyl alcohol (BA)/benzyl benzoate (BB) with diphenyl ether (BABB-D4) and BB-PEG-MA500, respectively, resulted in increased tissue autofluorescence in both the 488-nm and 561-nm imaging channels (Figure S1).

### The Effect of Temperature on Tissue Transparency and Fluorescent Signals

Temperature is a critical yet often overlooked factor influencing the optical transparency of biological samples. Although well recognized in physical systems, its impact on fluorescence imaging of tissue-cleared specimens is neglected. While tissue-clearing relies primarily on delipidation, most clearing techniques can only partially delipidate a specimen; residual lipids invariably remain within cellular membranes and lipid-rich structures. Since lipids are a major source of light-scattering in biological tissues, their physical state strongly influences optical clarity and consequently the ability to image fluorescent signals with high contrast at depth (Figure 2A). Since lipids scatter more light in the solid phase than in the liquid or disordered phase, we hypothesized that temperature-dependent phase transitions of residual lipids could significantly alter the transparency of biological samples.

**Figure 2.**
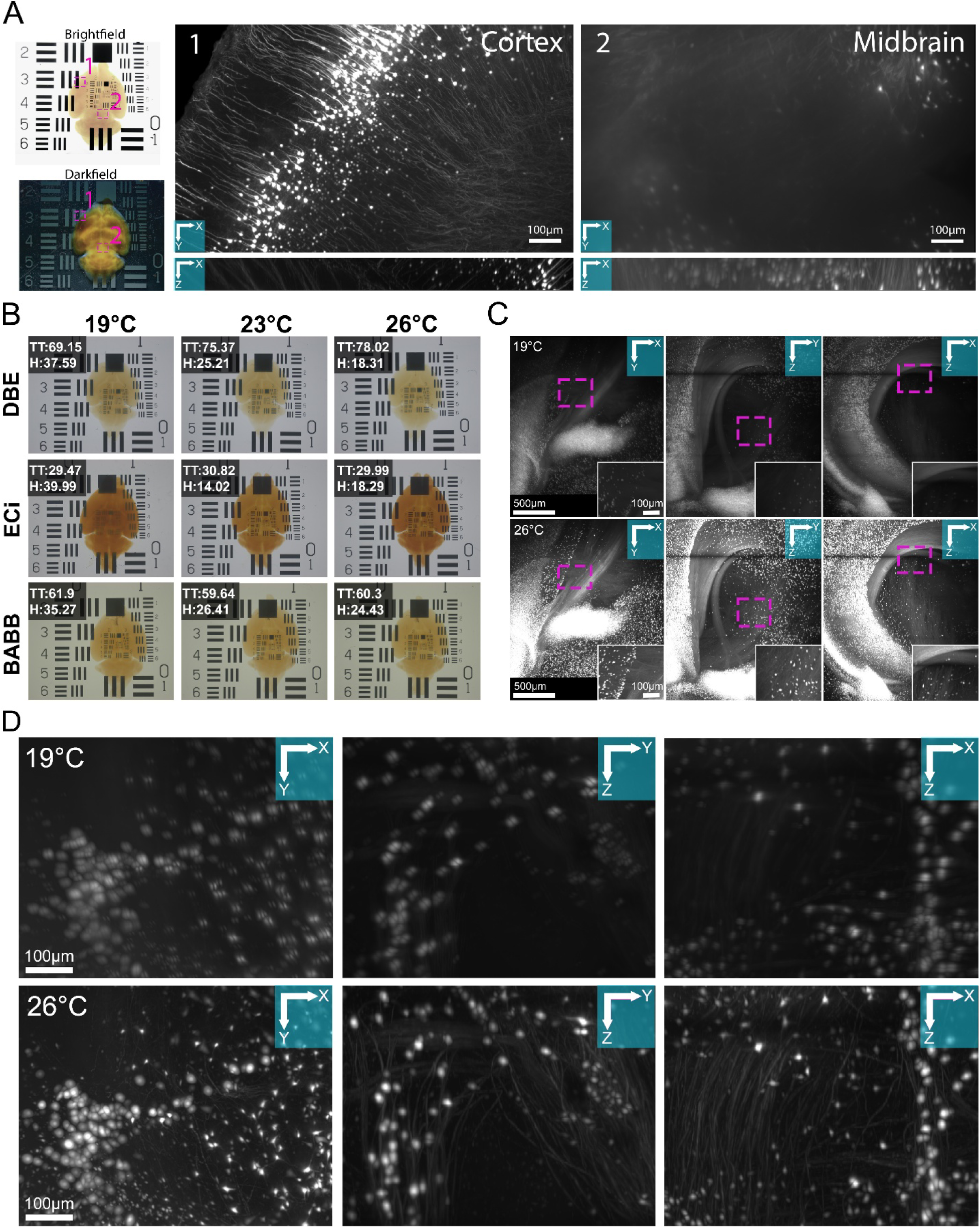
Impact of Temperature on Tissue Transparency. (A) Brightfield and darkfield images of a tissue-cleared Thy1-EYFP-H mouse brain on USAF chart. Dashed rectangles indicate regions of high (1) and low (2) transparency. Corresponding light-sheet images illustrate the effect of tissue transparency on fluorescent signal resolution. (B) Effect of temperature and refractive index (RI)-matching medium on TT and H, using mouse brains. RI-matching media: dibenzyl ether (DBE), ethyl cinnamate (ECi), and benzyl alcohol/benzyl benzoate (1:2; BABB). (C, D) Impact of temperature on fluorescent signal resolution, via light-sheet images of tissue-cleared Thy1-EYFP-H mouse brain, acquired with 4× NA-0.35 (C) and 12× NA-0.53 (D) objectives.

To systematically examine the impact of temperature on image quality, we first designed a custom temperature-controlled imaging chamber comprising a cuvette mounted on a heat sink connected to a digital temperature controller, allowing precise and stable thermal regulation during imaging (Figures S2A,B). We then integrated this imaging chamber into multiple imaging platforms, including a stereomicroscope, hazemeter, and mesoSPIM^35^ light-sheet microscope, to enable assessment of temperature-dependent changes in transparency, with transparency quantified as TT, H, and the occurrence of autofluorescence (Figures S2C–E). Next, we used our integrated imaging system to evaluate the influence of temperature on transparency in the imaging of cleared mouse brains, using a range of commonly used dehydration agents (MeOH, EtOH, 1-propanol, tB, and THF) and RI-matching media (ethyl cinnamate (ECi), DBE, and BABB).

Results showed an overall increase in TT with rising temperature, in the order of EtOH <MeOH< THF< tB< 1-propanol (Figures S2F,G). Among the RI-matching media, ECi RI-matched samples had the darkest coloration and consistently exhibited the lowest TT, whereas DBE and BABB demonstrated superior and temperature-dependent TT (Figures 2B and S2F,G). Next, we analyzed H across the same dehydration agents and RI-matching media, again using cleared mouse brain samples (Figures S2H,I). Haze decreased with increasing temperature under several conditions. Among the dehydration agents, MeOH and EtOH produced highest H values and showed strongest light scattering in darkfield, suggesting greater residual lipids. In contrast, tB, THF, and particularly 1-propanol, led to significantly lower H, corresponding to improved transparency (Figure S2I). Statistical comparisons revealed decreasing H across dehydration agents in the order of MeOH>EtOH>THF>tB>1-propanol (Figure S2I). When comparing RI-matching media, ECi consistently exhibited lower H than DBE and BABB, with minimal temperature dependence, whereas DBE and BABB showed reduced H and improved stability in optical performance with increasing temperature (Figure S2I). The overall decrease in H with increasing temperature for most RI-matching media suggests that elevated temperature reduces light scattering, potentially by inducing phase transitions in residual lipids, shifting them from an ordered to a disordered state. Ordered lipid domains at lower temperatures enhance RI heterogeneity and scattering, which limits transparency and imaging depth in cleared tissues. By reducing these heterogeneities, higher temperatures improve RI homogeneity throughout the sample, thereby increasing TT, reducing H, and ultimately enhancing volumetric image quality.

Heatmap visualization (see Methods) showed that spectral absorption decreased as temperature increased from 19°C to 26°C (Figure S2J), suggesting, as in the relationship between H and temperature, that increases in temperature reduce light scattering by promoting the transition of residual lipids from a more ordered to a disordered phase, thereby enhancing RI uniformity and overall optical transparency. Among the RI-matching media, DBE and BABB exhibited the lowest spectral absorption, whereas ECi maintained higher absorption and greater temperature stability (Figure S2J).

### Effect of Dehydration Agent, RI-matching Media, and Temperature on Autofluorescence

We next examined the effect of dehydration agents, RI-matching media, and temperature on autofluorescence in brain tissue from wild-type adult mice (C57BL/6J). Samples were processed using the same dehydration agents and RI-matching media as used above and then imaged at 488 nm and 568 nm at 19°C, 23°C, and 26°C (Figure S2K). Auto-fluorescence intensities were quantified and visualized as heatmaps (see Methods) for direct comparison across conditions (Figure S2L). Among the RI-matching media, DBE demonstrated superior optical stability for fluorescence imaging. Brain tissues in DBE exhibited the lowest overall autofluorescence at 488 nm and 568 nm, and their autofluorescence intensity, unlike TT and H, was minimally affected by temperature across both channels. In contrast, ECi induced higher autofluorescence than DBE. ECi-cleared brain tissues displayed the darkest coloration and highest light absorption (Figures S2F,L). Among the dehydration agents, MeOH produced low autofluorescence in brain tissue, despite its high H in cleared brain samples, particularly at a higher temperature (Figures S2F,L). This finding suggests that residual lipids are likely not the primary source of autofluorescence in brain samples. However, when dehydrated with MeOH, EtOH, or 1-propanol at lower temperatures (19°C), brain samples showed minimal autofluorescence at 568 nm but strong autofluorescence at 488 nm. Among the RI media, BABB produced the highest autofluorescence at 488nm and 568 nm in the brain with each dehydration approach and showed some degree of temperature influence (Figure S2K). In general, we also observed a modest change in autofluorescence with increasing temperature—either a slight decrease or increase depending on the dehydration agent and RI-matching medium. These results highlight that the RI-matching medium is the primary determinant of autofluorescence, while the dehydration agent and temperature represent additional parameters to modulate background signal.

Finally, to directly visualize temperature-induced improvements in image quality, we used our temperature-controlled chamber integrated with the mesoSPIM-ultra (described below) to acquire fluorescence images of sDBE-cleared Thy1-YFP-H brains at 19°C and 26°C using 4× (numerical aperture (NA=0.35) and 12× (NA=0.53) objectives (Figures 2C,D). The resulting images clearly demonstrate improved resolution and signal uniformity at elevated temperature, further indicating that controlled heating enhances optical transparency and imaging performance in tissue-cleared samples.

### The Effect of Sample Coloration on Fluorescence Imaging

It is well known that tissue-cleared samples often develop varying degrees of yellow-to-brown coloration during processing or later after RI matching (Figure 3A). To investigate how coloration affects TT and H, we recreated these effects in a controlled laboratory setting by using agar gels that differ in agar concentration, with Rosco optical filters of increasing color density placed separately in front of each agar sample (Figure 3B). Both TT and H values progressively decreased as the filters shifted from yellow to brown (Figures 3C,D). Increasing the agar concentration further reduced both parameters (Figures 3C,D), consistent with the phenomenon of a higher optical density leading to reduced scattering.

**Figure 3.**
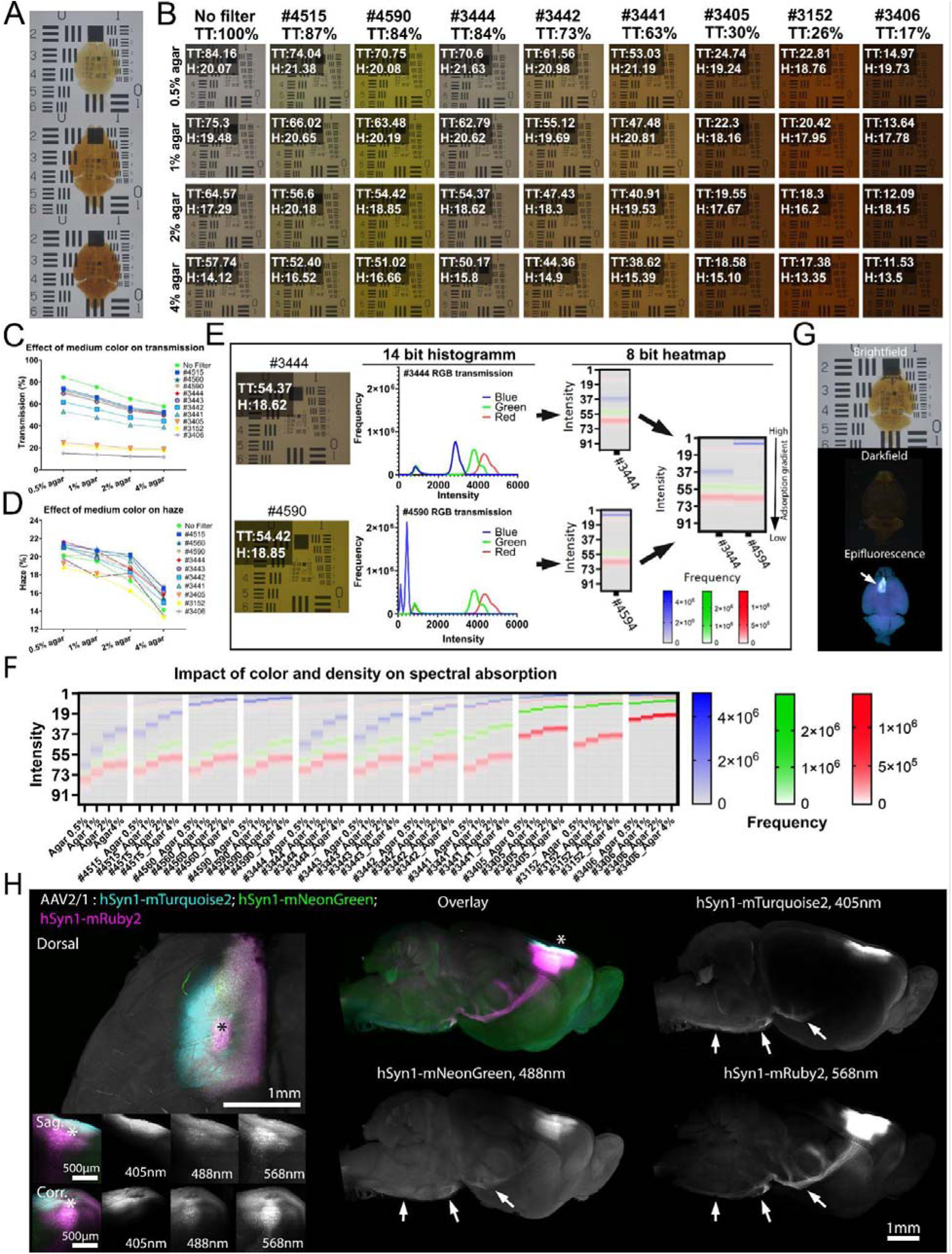
Impact of Coloration on Haze, Total Transmission, and Imaging Depth. (A) Representative brightfield images showing discoloration that develops in mouse brains following prolonged incubation in different RI-matching media. (B-D) The impact of sample coloration and medium density on TT and H, modeled using Rosco filters and agar gels of increasing concentrations (B). Manufacturer-provided TT values are indicated below each filter. Corresponding effects on TT and H are plotted in (C) and (D), respectively (n = 5 times measured). (E) Impact of coloration on spectral absorption (left; two filters that differ in color but result in similar TT and H), shown via red/green/blue (RGB) 14 bit histogram (middle), and the RGB histogram transformed as 8 bit heatmaps (right). (F) Impact of difference in coloration and agar concentration on spectral absorption, visualized as heatmaps derived from RGB histograms as described in (E). (G) Brightfield, darkfield, and epifluorescence images (top to bottom) of a cleared mouse brain labeled with an equal mixture of three AAV vectors expressing fluorophores excited at 405, 488, 568 nm. Arrow indicates injection site. (H) Showing brain from (G). Increased coloration preferentially attenuates transmission of shorter wavelengths. Multiview light-sheet images of mouse brain reveal reduced transmission at 405 and 488 nm, whereas signal transmission at 568 nm is largely preserved. Asterisks indicate the injection site; arrows indicate neuronal projections on the ventral side. Grayscale panels show individual imaging channels.

To directly assess the impact of coloration on spectral absorption, we compared two filters that exhibit similar TT and H but differ in color hue (Figure 3E). Corresponding heatmaps (see Methods) revealed that spectral absorption differed depending on the filter color, with greatest attenuation in the blue channel, moderate in the green, and minimal in the red (Figure 3F). These findings indicate that coloration selectively absorbs shorter wavelengths, while transmission of longer wavelengths remains largely unaffected (Figure 3F). Importantly, this also demonstrates that samples with similar TT and H values can differ markedly in coloration, and that coloration has an impact on fluorescence imaging performance that is distinct from that of TT or H. To validate these observations *in vivo*, we injected the motor cortex of C57BL/6J mice with an equal mixture (1:1:1) of three AAV vectors expressing three different fluorophores (mTurquois2, mNeon-Green and mRuby2) (Figure 3G). We then tissue-cleared the brains using our modified sDISCO protocol and incubated them for two months at room temperature in a light-unprotected environment using sDBE. Following this extended incubation period, a slight yellow coloration developed in the otherwise highly transparent brain tissue (Figure 3G). We illuminated the brain from both sides and successfully visualized the XFP signals from all three viruses. These signals were clearly visible at the dorsal injection site and in the ventral axonal projections located in the hypothalamus, pons, and medulla (Figure 3H). However, the axonal projections in the midbrain and thalamus showed variable visibility depending on the excitation wavelength. Consistent with the filter-based experiments, fluorescence imaging revealed that mTurquoise2 signals (excitation 405 nm) were the weakest and most attenuated, mNeonGreen (488 nm) showed intermediate brightness, and mRuby2 (568 nm) exhibited the strongest fluorescence intensity (Figure 3H). Together, these results demonstrate that sample coloration preferentially reduces transmission of shorter wavelengths, thereby disproportionately affecting blue and green fluorophores while sparing red-emitting ones. Moreover, our results highlight that coloration has a key impact on fluorescence-signal quality in cleared tissues.

### Effect of Dehydration Agent on Sample Morphology and Size During Tissue-Clearing

Dehydration-based tissue-clearing methods induce sample shrinkage. This can be advantageous, as smaller sample size reduces imaging time and data volume, and requires objectives with lower working distance. However, rapid or excessive dehydration can cause tissue deformation. We hypothesized that dehydration-agent polarity influences the rate of dehydration and, consequently, the degree of volume reduction (Figure 4A). To test this, we performed sequential dehydration of 4% PFA fixed mouse brain samples using the same five dehydration agents (MeOH, EtOH, 1-Propanol, tB, THF) and then quantified volume changes using a pycnometer (Figure 4B). Our results revealed variability in volume reduction during the initial dehydration steps. However, at 100% solvent concentration, the final sample volume was strongly correlated with dehydration-agent polarity (Figures 4A,B). Notably, volume reduction was lowest using MeOH and greatest using tB and THF, indicating that lower-polarity solvents promote greater tissue shrinkage under similar dehydration conditions.

**Figure 4.**
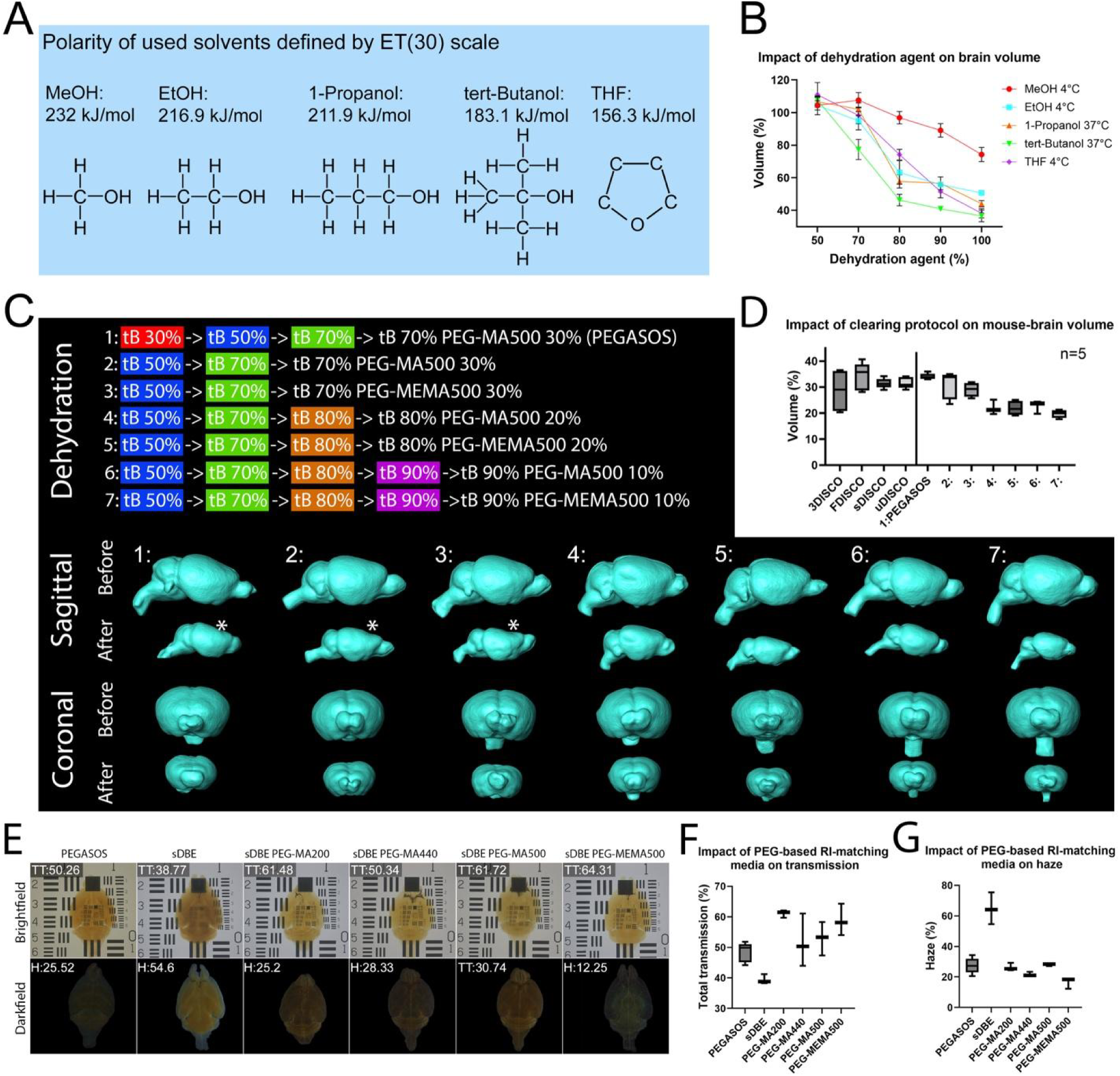
Effect of Dehydration Agent on Sample Morphology and Size during Tissue-Clearing. (A) Chemical structures and relative polarities (in kJ/mol) of commonly used dehydration agents. MeOH: methanol; EtOH: ethanol; THF: tetrahydrofuran. (B) Effect of sequential dehydration with different agents on mouse-brain volume, quantified via pycnometer. (n = 5 brains/condition). (C) Impact of different PEG-MEMA500-based dehydration protocols (top) on mouse-brain size and morphology visualized by 3D point cloud reconstruction. Sagittal (middle) and coronal (bottom) views are shown before and after tissue clearing. Asterisk indicates anisotropic cortical deformation. tB: tert-Butanol; PEG-MEMA500: polyethylene glycol methyl ether methacrylate 500. (D) Impact of different dehydration-based tissue-clearing methods on final volume of mouse brains, including the conditions shown in (C) (n = 5 brains). (E-G) Impact of different PEG-based RI-matching media on transparency, TT, and H using mouse brains. Representative brightfield and darkfield images are shown in (E), with corresponding TT and H measurements in (F) and (G), respectively (n = 3 brains). sDBE formulated with either PEG-MA or PEG-MEMA produced the highest TT and lowest H values. sDBE: stabilized dibenzyl ether; PEG-MA200: polyethylene glycol methacrylate 200; PEG-MA400: polyethylene glycol methacrylate 440.

### Optimization of tB-based Dehydration Using PEG-MEMA Additives

Dehydration using a tB gradient supplemented with polyethylene glycol methacrylate (PEG-MA) has recently gained attention for tissue-clearing, as part of the PEGASOS protocol^28^. The inclusion of PEG-MA facilitates efficient clearing while maintaining partial water retention and compatibility with hydrophobic RI-matching media, thus improving preservation of XFP fluorescence. Given that tB has one of the best optical performance profiles in our assays, i.e., the effect of dehydration-agent polarity on sample volume reduction (Figures 4A,B), and on TT and H (Figure S2F-I), we evaluated the combination of tB and PEG-MA as dehydration agents for mouse-brain samples, following the PEGASOS approach. First, we compared several variants of sequential dehydration using tB, followed by a final incubation in mixtures of tB:PEG-MA500 at 7:3, 8:2, and 9:1 ratios to test its impact on brain morphology and final sample size (Figure 4C). To assess morphological changes, we used a 3D-scanner-based point-cloud reconstruction together with a pycnometer-based volume analysis. We found that the 70:30 tB:PEG-MA500 mixture introduced morphological artifacts in the cortex while the 80:20 and 90:10 mixtures achieved isotropic shrinkage and greater reduction in tissue volume during dehydration (Figure 4C,D).

One limitation of the PEG-MA500-based PEGASOS protocol is optical distortion due to the highly viscous and non-uniform RI-matching medium, which also complicates sample handling. Further, the high melting point of PEG-MA (12-20°C) restricts its use in combination with dehydration agents such as THF, which can induce XFP quenching at these temperatures^24^. To overcome these limitations, we tested different PEG-based additives and found poly-ethylene glycol methyl ether methacrylate (PEG-MEMA500) to be a better alternative. PEG-MEMA500 has a relatively low viscosity and lower melting point (-1°C to 2°C), making it usable for dehydration agents where the incubation steps are performed at 4°C, such as THF.

Chemically, PEG-MA500 is hydroxyl-terminated and PEG-MEMA500 is methoxy-capped. Consequently, PEG-MA500 is more hydrophilic than PEG-MEMA500, allowing it to retain hydration and preserve XFPs^36,37^, while the more hydrophobic nature of PEG-MEMA500 offers better solvent miscibility while only modestly reducing hydration capacity. We tested multiple tB:PEG-MEMA500 formulations on soft tissue and found that gradual dehydration in tB up to 90%, followed by transfer to a 90:10 tB:PEG-MEMA500 mixture, produced maximum volume reduction without noticeable sample deformation (Figures 4C,D). To determine the best RI-matching medium, we dehydrated mouse brains for two days at 37°C in increasing tB concentrations (day 1: 50tB and 70tB; day 2: 80tB and 90tB) together with sDBE and different PEG-sDBE mixtures (all adjusted to an RI of 1.543) (Figure 4E). Hazemeter analysis showed that sDBE–PEG-MEMA500 exhibited the best overall performance, delivering highest TT, lowest H, and excellent sample integrity (Figures 4F,G).

To test the capacity of PEG-MEMA500 to clear bone and soft tissue and to be combined with THF, we developed a protocol that can be used at 4°C (Figure S3A). To test its efficacy, we performed whole-body tissue-clearing and vasculature imaging of adult casper zebrafish and brain vasculature of African turquoise killifish (ATK). Using zebrafish up to 20 months of age and ATK up to 5 months of age, we performed transcardial perfusion with a mixture of 10kDa dextran AF555nm and 10kDa dextran AF647nm. After fixation, the animals were incubated overnight in chilled acetone to remove some of the pigments and to reduce coloration of the samples after RI matching. Further, in this case and in our previous experience^4^, most of the melanin-based pigments could not be removed with this step. Therefore, after tissue-clearing (Figure S3B), we removed the melanin-rich eyes of the animals and used an Optimized Meso-Aspheric (OMA)-based LSFM^38^ to perform high-resolution imaging of fish vasculature within the intact heads, using a 16x objective (Leica; NA=0.6) (Figure S3C), and performed low-resolution whole-body imaging of zebrafish using a custom corrected 2x objective (NA=0.14) (Figure S3D). This allowed vascular reconstruction from organism-wide overviews down to fine intracranial vessels within the same sample, thereby providing a scalable platform for multiscale imaging of vertebrate anatomy and connectivity.

Together, these results demonstrate that a PEG-MEMA500-based clearing strategy enables whole-body, high-resolution imaging while preserving dextran-based labeling to visualize the entire vasculature of vertebrates. Our results also demonstrate that both solvent polarity and polymer additives have major impacts on tissue morphology and optical performance during dehydration-based clearing. Lower-polarity solvents such as tB and THF produced greater and more uniform sample shrinkage, while polar solvents such as MeOH and EtOH were less effective (Figures 4B,C). When PEG-MEMA500 was combined with sDBE for RI matching following either tB–PEG-MEMA500 dehydration for soft tissue or THF–PEG-MEMA500 dehydration for soft and hard tissue, the samples exhibited high-degree of transparency and structural stability (Figures 4C and S3D).

### An Optimized Tissue-clearing Protocol for XFP Signal Preservation in Mouse Brain

By optimizing conditions for optical performance, achieving high TT, low H, minimal autofluorescence, and signal retention, we developed an optimized tissue-clearing pipeline suitable for complex structures like mouse brain (Figure 5A). This workflow integrated the most effective reagents identified in our comparative analyses: (1) the use of Solution-1 for tissue permeabilization and decoloration^4^, (2) tB as dehydration agent with small incrementation steps during dehydration (70%, 80%, 90% tB) in combination with, (3) a 90:10 tB:PEG-MEMA500 mixture as a final dehydration step for higher lipid removal and to ensure controlled tissue shrinkage without causing tissue deformation, and (4) a 90:10 sDBE:PEG-MEMA500 mixture as the RI–matching medium to achieve maximum transparency while preserving fluorescence. In addition, we incorporated 3% Quadrol, a weak base known to raise pH and enhance XFP stability during clearing^10^. We evaluated protocol performance using brains from Thy1-EGFP-M mice with sparse neuronal labeling and ACTB-tdTomato mice with ubiquitous expression, monitoring histogram changes and quantifying fluorescence preservation at each clearing step (Figures 5A-E). We observed a modest decrease in fluorescence intensity at the 70% tB/3% Quadrol stage, followed by a partial recovery in the 90% tB/10% PEG-MEMA500 step. Importantly, both EGFP and tdTomato signals were preserved following the final 90% sDBE/10% PEG-MEMA500 RI–matching step, demonstrating retained fluorophore stability. When compared to the other clearing strategies tested, this protocol showed superior fluorescence preservation for both EGFP (mean: 0.001-fold; max: 0.25-fold), and tdTomato (mean: 0.001-fold; max: 0.22-fold). Hazemeter analysis of Thy1-eGFP-M brains revealed high TT and low H (Figure 5F). To test the compatibility of this workflow with exogenous dyes, we intracardially injected 10 kDa dextran AF555 into Thy1:GFP-M mice. Tissue-clearing and subsequent light-sheet imaging allowed visualization of the cerebral vasculature by AF555 along with EGFP-labeled neurons (Figure 5G).

**Figure 5.**
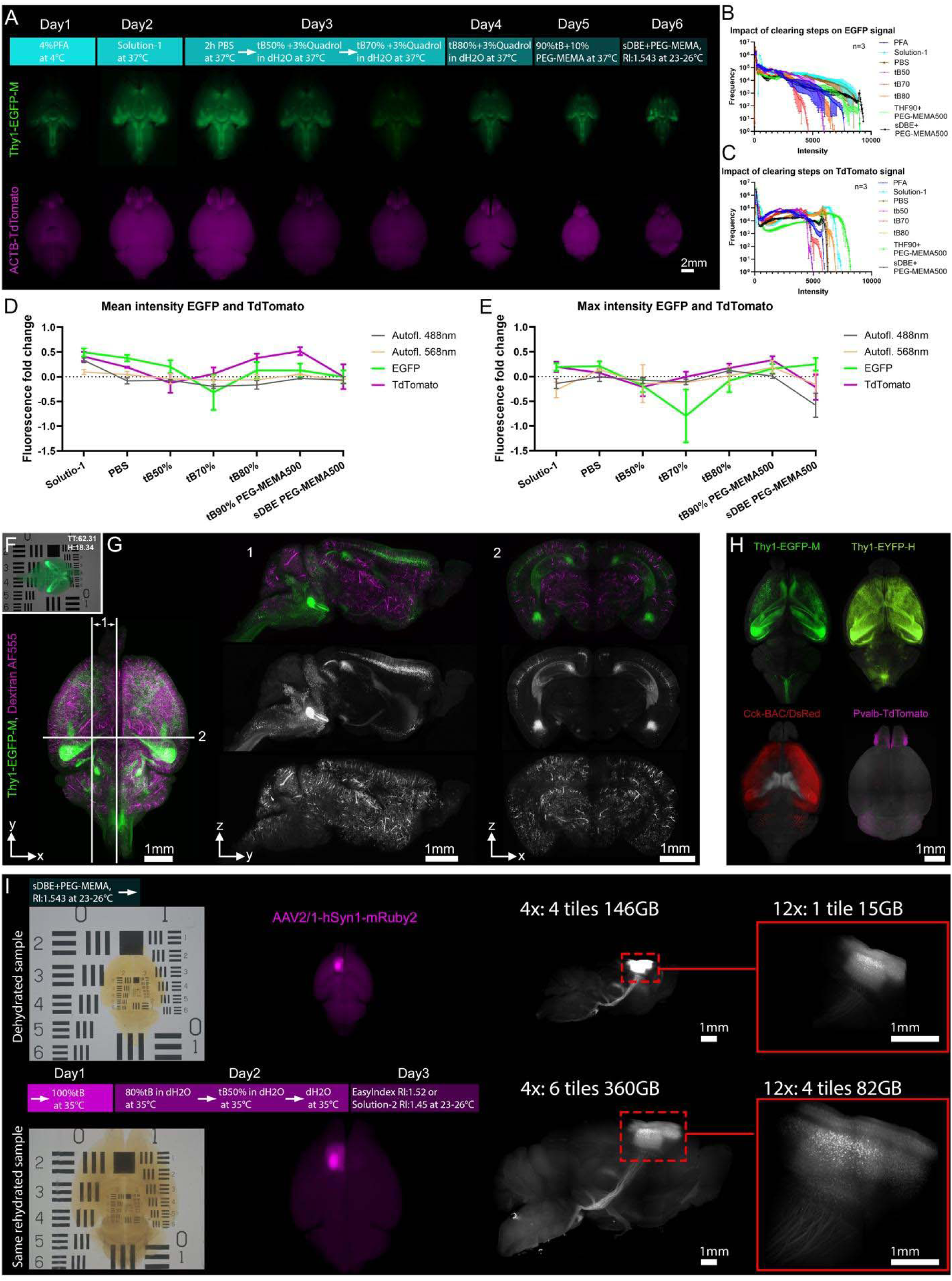
An Optimized Tissue-clearing Protocol for Robust XFP Signal Preservation in Mouse Brain. (A) Overview of the optimized tissue-clearing workflow on mouse brain, with representative epifluorescence images showing EGFP (green) and TdTomato (magenta) fluorescence retention at each step. PFA: paraformaldehyde; tB: tert-Butanol; PEG-MEMA500: polyethylene glycol methyl ether methacrylate 500. (B, C) Changes in EGFP (B) and tdTomato (C) fluorescence intensity throughout the tissue-clearing protocol, shown as intensity histograms (n = 3 brains). (D, E) Changes in mean and maximum autofluorescence and transgenic fluorescent signal across protocol steps, normalized to PFA-fixed baseline levels (n = 3 brains). (F) Cleared Thy1-EGFP-M mouse brain with our optimized protocol, demonstrating high TT, low H, and preserved epifluorescence. (G) Light-sheet images of a cleared Thy1-EGFP-M mouse brain after transcardial injection of 10-kDa dextran AF555nm, showing preserved transgene fluorescence (green) and brain-wide vascular labeling (magenta). Lines 1 and 2 indicate the planes of the optical sections. Grayscale images show individual channels. (H) Light-sheet images of cleared mouse brains expressing Thy1-EGFP-M (top left), Thy1-EYFP-H (top-right), Cck-BAC/DsRed (bottom left), or Pvalb-TdTomato (bottom right). (I) Comparison of dehydration- and water-based tissue-clearing on imaging data size and data quality, using a mouse brain injected with AAV2. Brightfield (left), epifluorescence (middle), and light-sheet images (right) of a brain cleared using a dehydration-based method (top). The same brain after rehydration and clearing with a water-based method (bottom). Insets show higher-magnification of dashed region; objective, tile number, and acquisition parameters are indicated above.

To further demonstrate versatility, we cleared CAG:GFP, CAG:tdTomato brain organoids and additional brains from Thy1-EGFP-M, Thy1-EYFP-H, Cck-BAC/DsRed, and Pvalb-tdTomato mice, all of which retained their native fluorophore signals—EGFP, EYFP, DsRed, and tdTomato, respectively (Figures 5H and S4). We observed high XFP signal-to-noise ratio (SNR) in brain organoids (Figure S4A) and across diverse brain regions such as the cerebellum (Figures S4B-F), cortex (Figures S4B,C,F), hippocampus (Figures S4B,F), olfactory bulbs (Figures S4B,C,F,G), pons, medulla, and spinal cord (Figure S4D) while maintaining cellular and subcellular resolution, as demonstrated by visualization of dendritic spines on cortical neurons (Figure S4E) and the dendritic arborization of cerebellar interneurons (Figure S4H, Video S1).

To test our protocol in different rodents and on structures sparsely expressing XFP, we performed rAAV2/5-hSyn-hChR2(H134R)-EYFP virus injections in adult Long-Evans rats to reveal ventral posteromedial nucleus (VPm) projections into the primary somatosensory cortex, as well as dentate gyrus (DG) projections to the CA2/3 regions of the hippocampus (Figure S4I). Further, AAV-hSyn-DIO-EGFP virus injections in *Crh*-Ires^Cre^ mice visualized the local axons and axon collaterals of CRH+ neurons together with long-range efferents to the ventrolateral thalamus, and the somatosensory and somatomotor cortices (Figure S4H). In sum, the incorporation of PEG-MEMA500 in dehydration and RI matching offers low viscosity and high solvent compatibility with minimal loss of fluorescence.

### Interchangeable Solvent- and Water-based Imaging

An additional goal of this work was to establish a reversible clearing workflow that permits transition between solvent- and water-based imaging modalities. Such an approach would enable rapid low-resolution screening of large specimens in a compact solvent-cleared state, followed by rehydration and higher-resolution imaging of selected regions after sample expansion. To test this, we first injected a mouse brain with an AAV2/1-hSyn1-mRuby2 virus, tissue-cleared it according to the protocol in Figure 5A and imaged the brain on a light-sheet microscope. We then rehydrated the brain, embedded it in 2% low-melting agarose and subsequently performed RI matching with EasyIndex (RI:1.52; LifeCanvas Technologies), which is suitable for aqueous-based clearing methods. The rehydrated brain was imaged on the same light-sheet system using identical optical settings. This interchange between a dehydration-based and water-based tissue-clearing approach allowed for imaging with fewer tiles to generate an overview stack to identify areas of interest and subsequently, after rehydration, to image the same areas with higher resolution due to physical expansion of the sample (Figure 5I).

### mesoSPIM-ultra Design Enables High-resolution Imaging of Tissue-cleared Samples

Neither the original mesoSPIM^35^ nor the more recently published benchtop mesoSPIM^39^ utilize detection objectives that are corrected for imaging in high-index clearing media. To overcome this limitation, we upgraded the mesoSPIM with: (1) clearing-optimized immersion objectives, improving lateral resolution two-fold and axial resolution four-fold; (2) a large-format CMOS camera (Teledyne Kinetix), increasing FoV by 36%; and (3) enhanced tile-stitching capabilities, reducing post-processing errors. These improvements enable cellular-to-subcellular imaging across larger fields with improved efficiency (Figures 6A,B).

**Figure 6.**
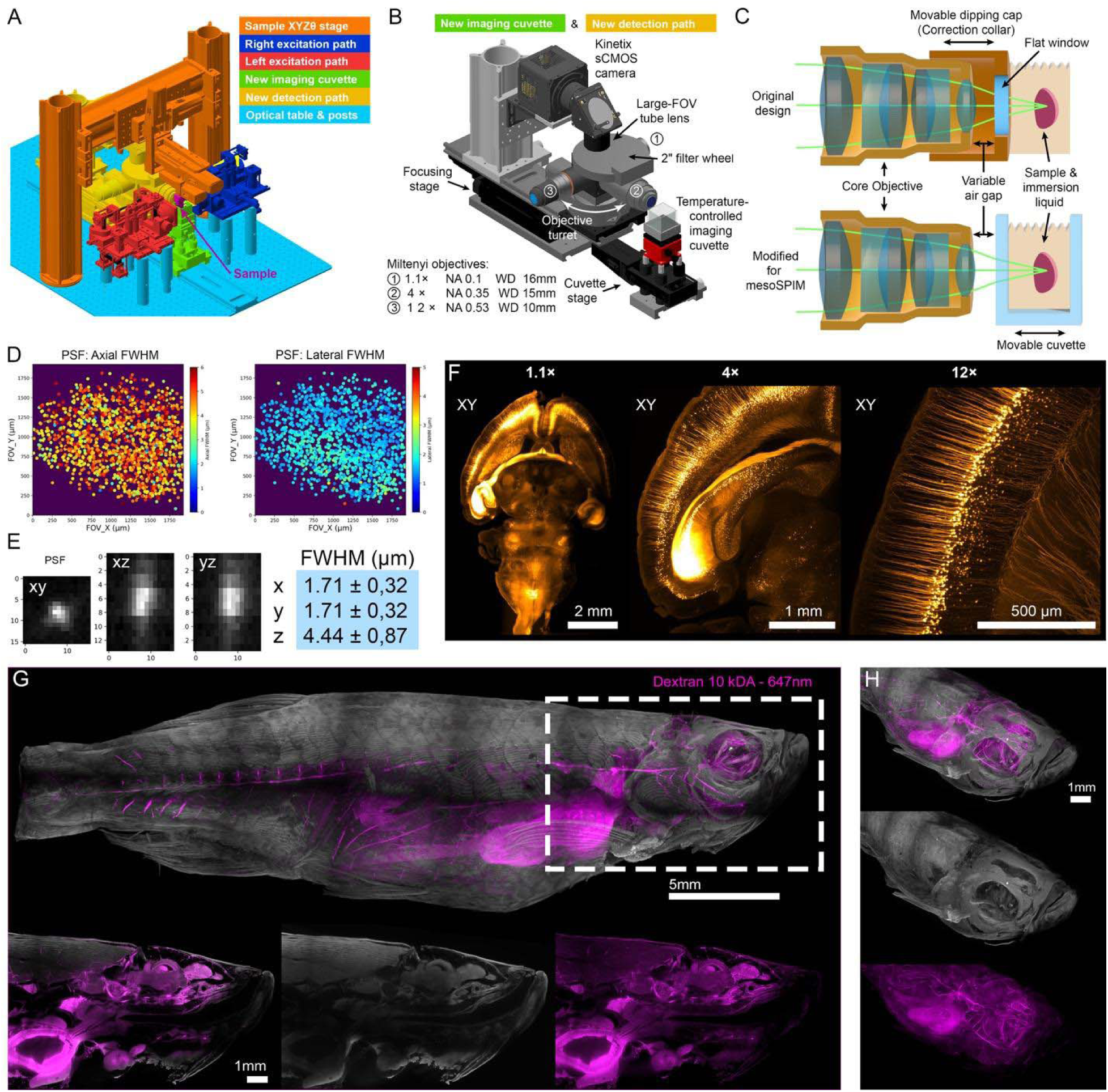
mesoSPIM-ultra Configuration Enhances Optical Performance and Imaging Capabilities. (A) Color-coded schematic of the mesoSPIM-ultra setup. (B) Overview of the new detection and cuvette components. Three objectives (1.1×, 4×, and 12×) are mounted on a horizontal objective turret. Both the detection path and cuvette are mounted on motorized linear stages. (C) Design and adaptation of Miltenyi Biotec immersion objectives for mesoSPIM-ultra. The original dipping objectives include a cap with a flat window and an adjustable air gap controlled (top) by a correction collar to minimize medium-induced spherical aberration. For mesoSPIM-ultra use, the dipping caps are removed, with the cuvette wall serving as the flat interface. Because the objectives are not parfocal and have different working distances, both the objective and cuvette positions are adjustable via dedicated focusing and cuvette stages. (D) Point spread function (PSF) maps axial (left) and lateral (right) full width at half maximum (FWHM) of fluorescent beads imaged with the 12× objective of mesoSPIM-ultra. (E) Representative PSFs and average full width at half maximum (FWHM) values from the dataset are shown. in (D). (F) Comparison of the field of view (FoV) achieved in a Thy1-YFP-H mouse brain using 1.1×, 4×, and 12× objectives. (G-H) Whole-body imaging of autofluorescence (gray) and 10-kDA dextran 555nm (magenta) labelled vasculature in adult zebrafish (20 months old) using a 4× NA-0.35 objective. Dashed area shows clipping plane of autofluorescence and vasculature (left), autofluorescence (middle) and vasculature (right).

We incorporated three immersion objectives that were designed by Miltenyi Biotec for the Ultramicroscope Blaze into our mesoSPIM-ultra: 1.1× (NA=0.1), 4× (NA=0.35), and 12× (NA=0.53) (Figure 6B). Each objective featured a correction collar for imaging in RI-matching media ranging from RI 1.33-1.56 (Figure 6C). Two major design challenges were associated with this modification. First, the original mesoSPIM used an Olympus MVX-10 Macro-Zoom microscope with continuous zoom (NA=0.25), which we replaced with a motorized objective turret capable of precisely positioning the heavy objectives (e.g., the 4× weighs 1.62 kg) (Figure 6B). Second, the Miltenyi Biotec objectives were designed for a vertical detection path whereas the mesoSPIM utilizes a horizontal detection path with air objectives. These dipping lenses minimize spherical aberration by adjusting the relative optical path lengths in air and immersion medium via movement of the front window. In the mesoSPIM, switching objectives would normally require breaking liquid contact, complicating automation and increasing leakage risk. Therefore, we removed the dipping caps (Figure 6C) and used the side wall of an immersion cuvette as the optical interface. The objectives therefore never encountered the immersion medium directly. To emulate correction-collar functionality, we mounted the cuvette on a motorized stage along the optical axis, allowing computer-controlled adjustments of the optical path lengths in air and liquid (Figure 6C). The modified mesoSPIM-control software automatically adjusts cuvette positioning when objectives are switched, based on user-defined calibration setpoints. Switching between objectives is facilitated by a horizontal heavy-duty rotation stage (PI MICOS L-611) capable of holding as many as four objectives. Due to the large back apertures of the new objectives, we also incorporated a two-inch emission filter wheel (ZWO 7×2”).

Using the mesoSPIM-ultra, we first evaluated the point spread function (PSF) (Figures 6D,E). The modified setup provided working distances of 17mm, 16mm, and 11mm for the 1.1×, 4×, and 12× objectives, respectively. At 12× magnification, the measured full width at half maximum (FWHM) was 1.71±0.32 μm in the x-y plane and 4.44±0.87 μm along the z-axis (Figure 6E), confirming high axial and lateral resolution suitable for mesoscale imaging. To assess optical performance in cleared tissue, we cleared Thy1-YFP-H mouse brains and imaged them using the 1.1×, 4×, and 12× objectives. The redesigned optical system yielded expanded FoVs of 20.8mm², 5.6mm², and 1.92mm² for the 1.1×, 4×, and 12× objectives, respectively (Figure 6F), with the possibility of whole-body imaging of samples such as adult zebrafish over three cm in length (Figures 6G,H, Video S2). Together, these modifications significantly enhanced imaging resolution, FoV, and optical correction in high-RI media, extending the mesoSPIM platform’s capability for high-quality volumetric imaging of tissue-cleared samples.

### Combination of Improved Tissue-clearing Protocol and mesoSPIM-ultra Enables Single-neuron Tracing of a Multi-cm-scale Projectome

Clozapine N-oxide (CNO)-inducible activation of an AAV-based DIO-DREADD system in *Chx10^Cre^* mice enables selective stimulation of *Chx10^Cre^*-expressing gigantocellular (Gi) reticulospinal neurons, which drive ipsilateral head and limb orientation and induce ipsilateral turning behavior in mice^40^. While previous efforts to map the projections of these neurons using conventional histological sectioning revealed that *Chx10^Cre^*Gi neurons innervate multiple targets within the brainstem and spinal cord^40,41^, the long-range trajectories, complex branching patterns, and multi-centimeter organization of this circuit precluded its *en block* visualization due to inherent limitations to serial tissue sectioning.

To demonstrate the combined power of our physicochemically optimized tissue-clearing protocol and mesoSPIM-ultra imaging, we reconstructed the multi-centimeter projection architecture of *Chx10^Cre^*Gi neurons. We first injected *Chx10^Cre^* mice (*n*=12) with an AAV vector carrying a loxP-flanked DREADD (hM3Dq) construct, which allowed CNO-dependent neuronal activation (Figures 7A,B) and simultaneously genetic labeling of these neurons (Figure 7C). Initially, we quantified the number of AAV-labeled Gi reticulospinal neurons using the syGlass VR platform. Approximately ≥5,000 Gi neurons were counted, providing a population estimate for mice (Figures 7D-F). To achieve long-range single-neuron reconstruction, we cleared the entire CNS (brain plus spinal cord) of *Chx10^+^* mice (*n*=2) and imaged the samples at 26°C using 12× magnification to obtain high-resolution 3D datasets spanning the entire rostrocaudal axis of the CNS (Figures 7C and S5A-C). Using the syGlass VR platform, we performed single-neuron tracing across the entire specimen, thus revealing long-range reticulospinal projections at single-neuron resolution (Figures 7G-K, Video S3). These reconstructions had projection motifs that were not captured using conventional section-based approaches^40,41^, including previously underappreciated cranial motor, ascending, contralateral, inferior olive, and multi-segment spinal pathways. These results indicate that *Chx10*^Cre^ Gi neurons comprise a projectionally-diverse reticulospinal population, with individual neurons exhibiting collateral architectures (Figures 7G-K) that could coordinate ipsilateral orienting movements through distributed projections in multiple spinal-cord segments.

**Figure 7.**
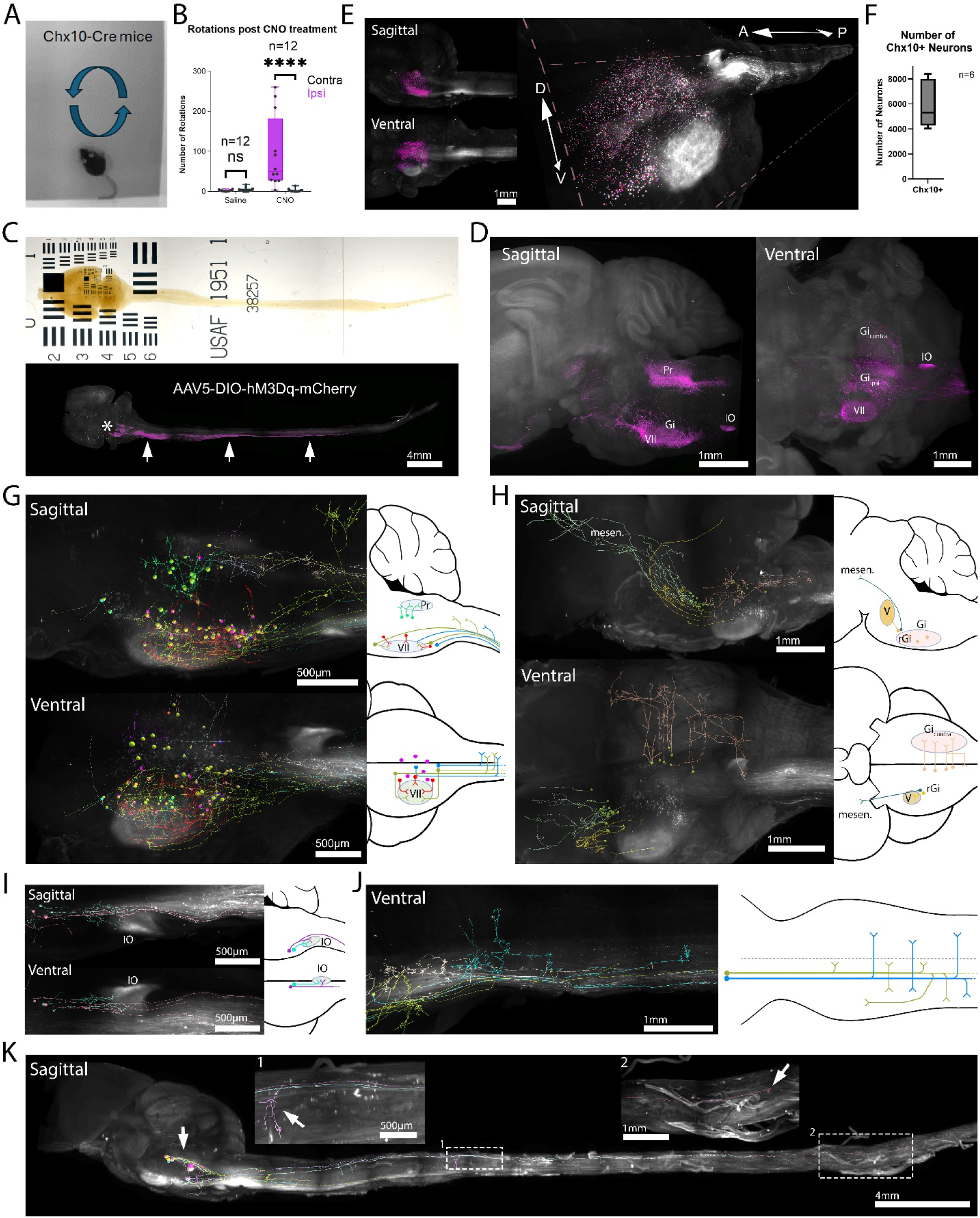
Integration of Behavioral and Motor Projectome Analysis in *Chx10^Cre^* Mice. (A, B) Phenotypic characterization of *Chx10^Cre^* mice after injection of Cre-dependent hM3Dq-DREADDs into the left gigantocellular nucleus (Gi), and subsequent administration of clozapine-N-oxide. Arrows in (A) indicate ipsilateral turning behavior, and (B) shows the significantly increased number of ipsilateral rotations in these mice. (C-K) Brain or both brain and spinal cord together from *Chx10*^Cre^ mice (A) were harvested for subsequent processing and analysis. (C) Top: Brightfield image of a tissue-cleared *Chx10^Cre^* mouse brain and spinal cord. Bottom: corresponding light-sheet image showing viral injection sites (asterisk) and *Chx10* neuronal projections (arrows). (D) Light-sheet image of the viral injection region. (E) Light-sheet images showing manual annotation of *Chx10* neuronal cell bodies (magenta) using syGlass VR software, with quantification. (F) Six *Chx10^Cre^*mice from (A) with robust turning behaviors were quantified for number of labeled neuronal cell bodies. (G-K) Light-sheet images showing manual tracing of *Chx10* neuronal projection areas and arborization patterns using syGlass VR, with corresponding projection schematics. (G) Gi neurons projecting locally into VII (red), into VII and the spinal cord (yellow), exclusively into the spinal cord (blue), or locally into Pr (green). (H) PnC neurons projecting into V (yellow), into the mesencephalon (green), and Gi_ipsi_ neurons projecting into the Gi_contra_ region (pastel pink). (I) Gi neurons projecting into the IO (cyan) or both the IO and the spinal cord (magenta). (J) Gi neuronal projections into the spinal cord showing multiple bifurcation points—either on one side of the midline (yellow) or on both sides (blue). The dotted line in the schematic drawing represents the midline. (K) Chx10 neuronal projections along the entire spinal cord. Notably many Chx10 Gi axons descend ipsilaterally and then branch contralaterally at the level of their target spinal segment revealing a prominent bilateral innervation pattern. Insets: (1) bifurcation points (arrow, inset 1) (2) terminal ends of projections (arrow, inset 2). VII: facial motor nucleus; IO: inferior olivary complex; Gi: gigantocellular reticular nucleus; rGi: rostral Gi; ipsi: ipsilateral to the injection site; contra: contralateral; Pr: prepositus nucleus, V: motor nucleus of the trigeminal nerve; mesen: mesencephalon.

## DISCUSSION

Tissue-clearing and light-sheet imaging have transformed 3D visualization of intact biological systems. As the number of clearing protocols has expanded, the challenge has shifted from technical feasibility toward rational method selection and optimization. Numerous aqueous- and solvent-based approaches have been developed, each balancing transparency, fluorescence preservation, tissue expansion or shrinkage, and imaging compatibility differently. Water-based methods^6–9,17,18,30,42^ emphasize hydrogel stabilization and XFP preservation, whereas solvent-based approaches^21–26,28,29,43^ prioritize rapid clearing, higher transparency and compatibility with both soft and mineralized tissues. However, mechanistic insights into these clearing strategies remain limited.

Rather than introducing an additional empirically optimized protocol, we established a principle-driven framework linking dehydration chemistry, RI matching, temperature, and tissue coloration to measurable optical outcomes. By integrating effective components from existing methods and systematically benchmarking their performance, we combined established strengths with quantitative optimization to generate a more versatile and mechanistically informed workflow.

Our findings demonstrate that tissue transparency cannot be adequately described by visual inspection alone, as it fails to distinguish absorption from scattering. To our knowledge, only one prior study employed a hazemeter for tissue analysis, measuring transmission alone and not haze. Further, the measurements in this study were performed outside the RI media, introducing surface-scattering artifacts^44^. By combining TT and H measurements in RI-matched tissues, we show that samples with similar apparent transparency can differ substantially in scattering behavior, demonstrating that transparency reflects distinct contributions of absorption and scattering governed by RI heterogeneity within tissue.

Our results further identify dehydration-agent polarity and solvent incrementation as major effectors of tissue morphology and optical quality. Consistent with prior studies^23,24,28,45^, lower-polarity solvents such as THF and tB achieve better transparency in lipid-rich tissues, likely because of more efficient lipid removal. We extend these observations by linking solvent polarity and dehydration-step transitions to tissue shrinkage, structural preservation, and RI heterogeneity, highlighting that dehydration chemistry influences morphology and residual lipid distribution, in addition to transparency.

A major conceptual advance emerging from this study is our identification of imaging temperature as a determinant of optical quality. Earlier work focused on elevated temperatures during delipidation^45^ or solvent-induced fluorescence loss^24^, whereas the effect of imaging temperature itself remained unexplored. Across multiple microscopy modalities, elevated temperatures improved transparency and reduced haze under several RI-matching conditions. These observations suggest that residual lipids undergo temperature-dependent phase transitions that improve RI homogeneity and reduce scattering. In particular, overly strong air conditioning in microscopy labs can negatively affect tissue transparency.

We also identified tissue coloration and choice of RI medium as independent variables influencing autofluorescence and fluorescence imaging quality. Yellow or brown discoloration occurs in most clearing approaches^7–9,23,24,26,28,45–48^, yet its optical consequences have remained uncharacterized. Our data demonstrate that coloration disproportionately attenuates shorter wavelengths, causing greater signal loss for blue and green fluorophores even in highly transparent tissues. These findings further highlight that transparency and haze measurements alone are insufficient predictors of fluorescence imaging performance. Building on these insights, we optimized a tB-based dehydration workflow supplemented with PEG-derived additives. PEGASOS previously demonstrated that PEG-containing solvents improve fluorescent-protein preservation in solvent-based clearing^28^. Expanding this concept, we show that PEG-MEMA500 improves compatibility with delipidation temperature for various dehydration agents, while the combination of PEG-MEMA500 with sDBE as an RI medium preserves XFP fluorescence, increases SNR, and maintains favorable imaging properties.

To fully leverage these chemical advances, we redesigned the mesoSPIM platform to complement the optical properties of cleared tissues. By integrating modified immersion optics, a large-chip sCMOS camera, temperature-controlled imaging, and automated optical-path correction, we improved high-resolution imaging across centimeter-scale sized specimens. We further developed an automated strategy to minimize spherical aberrations by translating the immersion cuvette along the detection axis to the optimal position for each objective, replacing manual adjustment of objective correction collars. This design enables rapid objective switching without fluid spills or custom imaging chambers, improving usability while reinforcing the principle that tissue-clearing and microscope engineering should be co-optimized.

The biological application presented here illustrates the broader potential of this integrated framework. Previous studies showed that *Chx10*^+^ Gi reticulospinal neurons contribute to orienting behaviors^40^, but conventional histology could not unequivocally resolve their local *vs.* long-range axon collaterals^40,49^. By combining optimized clearing, high-resolution mesoSPIM imaging, and syGlass-based virtual-reality tracing, we reconstructed multi-centimeter neuronal trajectories spanning the entire mouse CNS. These data resolved a central question, whether spinal-cord and inferior-olive projections arise from distinct neuronal populations or collateral branches of the same command neurons^40,41^. Our reconstructions provide substantial anatomical evidence that at least a subset of *Chx10^+^* Gi neurons collateralize to both the spinal cord and inferior olive, thus offering an anatomical substrate through which brainstem commands may be relayed to olivocerebellar circuits^50^. This application highlights the strength of our workflow in resolving long-range axonal trajectories and branching patterns across deep, axon-dense, and myelin-rich CNS regions.

In conclusion, our study establishes a mechanistic and quantitative framework for dehydration-based tissue-clearing that integrates optical physics, solvent chemistry, and microscope engineering into a unified imaging strategy. By systematically benchmarking transparency, haze, autofluorescence, morphology, and fluorescence preservation, we define principles for rational optimization and standardization of intact-tissue imaging workflows. More broadly, these findings shift tissue-clearing from largely protocol-driven empirical optimization toward predictive and customizable design, based on measurable physicochemical and optical parameters, enabling advances in volumetric imaging, developmental biology, and systems neuroscience.

## Limitations of the study

Optimal clearing conditions remain tissue-dependent, reflecting differences in lipid composition, extracellular-matrix organization, pigmentation, and endogenous autofluorescence across species, developmental stages and different organs. Large lipid-rich tissues may require modified detergents^51,52^ or elevated dehydration temperatures^53^, whereas tissues with lower lipid content can often be efficiently cleared under milder conditions. Solvent-based shrinkage, while advantageous for imaging throughput, may require calibration for quantitative morphometric analyses. Additionally, several results presented here are correlative and predictive in nature, derived from observed relationships between optical performance and the physical and chemical properties of clearing reagents and biological samples. While these relationships provide a useful framework for rational protocol design, mechanistic validation at molecular and ultrastructural levels will be required to establish causal relationships underlying the observed transparency changes. Future studies integrating ultrastructural analyses, lipidomics, and direct biophysical measurements will therefore be essential to fully resolve these mechanisms. Finally, although VR tracing proved highly effective for neuronal reconstruction, manual analysis may still introduce user-dependent bias. Moreover, the neuronal projections analyzed here were derived from mouse tissue, whereas corresponding spinal and supraspinal networks in larger mammals, including humans, span substantially greater distances and may present additional imaging, clearing, and reconstruction challenges. Future integration of adaptive optics, automated parameter selection, and standardized transparency metrics should further improve reproducibility and cross-laboratory comparability.

## RESOURCE AVAILABILITY

### Lead contact

Further information and requests for resources, reagents, and data should be directed to and will be fulfilled by the lead contact, Prayag Murawala.

### Materials availability

This study generated no new unique reagents. Viral constructs and plasmids used in this work are available from Addgene or VVF Zurich as specified in the Key Resources Table.

### Data and code availability

This study did not generate new code. All datasets supporting the findings of this work are available from the Lead Contact upon reasonable request.

## ACKNOWLEDGMENTS

This work was supported by NIH/NIGMS grants R35GM161492 to PM, and by COBRE grant P20GM104318 (PI: Iain Drummond), on which PM served as a Research Group Leader. FFV was supported by the Salsbury Cove Research Fund (SCRF, MDIBL) and a Branco Weiss Fellowship administered by ETH Zürich. OK was supported by Lundbeck Foundation Professorship R310-2019-21 and a Novo Nordisk Laureate Grant. JMC was supported by EMBO Long-Term Fellowship ALTF 421-2018 and Lundbeck Foundation Postdoctoral Fellowship R3472020-2393. TH was supported by ERC-2020-AdG-101021016. NP was supported by Elise Richter Fellowship FWF project V931-B. HUD was supported by FWF-P35614. We acknowledge the Light Microscopy Facility (LMF; RRID:SCR_019166) at MDIBL, supported by NIH/NIGMS COBRE grant P30GM154610. We thank Stephen Sampson for manuscript proofreading and Paul Paetzold and Maike Kreutz for assistance with the mesoSPIM-ultra setup.

## AUTHOR CONTRIBUTIONS

Conceptualization: MP, JMC, SS, AA, JAK, TK, IA, TH, MTJ, VK, OK, HUD, FFV, PM; Methodology: MP, SS, FFV; Formal analysis: MP, JMC, NP, PP, AMC, FFV; Resources: PeS, JH, EB, JAK, VK, HUD, OK, PM; Investigation: MP, JMC, SB, AA, JR, SCP, NP, MSM, KJ, MW, PA, PaS, SK, RH, MEK, DR, MM, JUH, TK, TH, IA, MTJ, FFV; Technical assistance: SB, SCP, RPS, MSM, KJ, PaS, MW, PP, AMC, RH, PA, MEK, JHG.; Supervision: HUD, FFV, PM; Writing and editing: MP, JMC, JAK, TK, TH, IA, MTJ, VK, OK, FFV, PM. All authors reviewed and approved the manuscript.

## DECLARATION OF GENERATIVE AI

ChatGPT was used for text editing. All outputs were reviewed and verified by the authors.

## STAR★METHODS

### EXPERIMENTAL MODEL AND SUBJECT DETAILS

#### Mice (Mus musculus)

##### Austria

Adult male and female C57BL/6J, *Crh*-Ires ^Cre^, and CckBAC/DsRed mice (2-6 months old) were bred and maintained under specific-pathogen-free (SPF) conditions at the Medical University of Vienna. Animals were housed in standard cages at 22 °C on a 12 h:12 h light-dark cycle with food pellets and water provided *ad libitum*. All experimental procedures complied with Austrian national ethical regulations and were approved by institutional authorities.

##### United States

Adult male and female C57BL/6J, 129S6, 29S6-Pkd1<RC>, STOCK Tg(Thy1-EGFP)MJrs/J, B6.Cg-Tg(Thy1-YFP)/HJrs/J (hemi), and C57BL/6-Tg(Pvalb-tdTomato)15Gfng/J (hemi) mice (2-10 months old) were obtained from The Jackson Laboratory (Bar Harbor, ME, USA; strain #000664 and derivatives). Mice were maintained under SPF conditions in pressurized individually ventilated (PIV) racks at 23 °C and 50 % humidity on a 12 h:12 h light–dark cycle, with acidified drinking water and LabDiet 5K52 chow (6 % fat) available *ad libitum*. All procedures were approved by the Institutional Animal Care and Use Committee (IACUC) of The Jackson Laboratory (protocols #23-059, #13003 and #14010).

##### Denmark

Adult male and female *Chx10*^Cre^ mice^40^, at least 8 weeks old and heterozygous for the Cre allele, were bred and maintained under SPF conditions at the University of Copenhagen. Animals were housed in standard cages at 23-24 °C and 45-50 % humidity on a 12 h:12 h light–dark cycle with *ad libitum* access to food and water. All procedures were approved by the Danish Animal Experiments Inspectorate (Dyreforsøgstilsynet) (permits 2017-15-0201-01172 and 2022-15-0201-01131) and reviewed by the University of Copenhagen Ethics Board (plans P19-134, P21-323, P22-502, A20-160, and A23-154).

#### Gerbil (Meriones unguiculatus)

Adult male and female gerbils (2-4 months old) were bred and maintained under SPF, air-conditioned conditions at 22 °C on a 12 h:12 h light-dark cycle with food and water *ad libitum*. All procedures were approved by the Animal Care Committee of Sachsen-Anhalt, Germany (permit #42502-2-1103 LIN MD) and performed in accordance with the NIH Guide for the Care and Use of Laboratory Animals (2011).

#### Rat (Rattus norvegicus)

Adult Long-Evans rats (2-4 months old; Charles River Laboratories) were housed under SPF, air-conditioned conditions at 22 °C with a 12 h:12 h light-dark cycle and free access to food and water. All procedures were approved by the Austrian Ministry of Science and the Medical University of Vienna (permit #66.009/0281-WF/V/3b/2015).

#### Zebrafish (*Danio rerio*)

Adult casper zebrafish (5–20 months old) were maintained at 28.5 °C in recirculating water (pH ≈ 7.2; conductivity 1000–1200 µS/cm) on a 14 h:10 h light–dark cycle. All experimental procedures were approved by the Mount Desert Island Biological Laboratory (IACUC protocol #AUP 20-02).

#### African Turquoise Killifish (*Nothobranchius furzeri*)

Adult African turquoise killifish (5 weeks old) were maintained at 28.5 °C in recirculating water (pH 7.4–7.6; conductivity ≈ 2700 µS/cm) on a 12 h:12 h light–dark cycle. All experiments were approved by the Mount Desert Island Biological Laboratory (IACUC protocol #AUP 21-03).

### METHOD DETAILS

#### Transcardial Perfusion and Sample Fixation

##### Rodents

Preparation of perfusion solutions: We mixed 100µl heparin (Sigma-Aldrich, H3393-250KU) in 500ml 1xPBS for initial blood removal. We then mixed 4% PFA into the above solution for fixation and maintained this solution ice-cold. For blood-vessel staining, we mixed a dye solution of 1mg/ml dextran 10kDa-555 and/or 0.5 mg/ml dextran 10kDa-647 in 0.8xPBS + 1% gelatin.

Perfusion procedure: Animals were anesthetized by intraperitoneal (IP) injection of 4% tribromoethanol 0.3ml/per 10g body weight. Anesthesia was confirmed by pinching reflexes of the paws. Mice were perfused transcardially with 50ml of PBS, while rats and gerbils were perfused transcardially with 200ml-500ml of PBS (RT-37°C). Initial quality of perfusion can be assessed by the liver turning completely white. This was followed by a 100-ml transcardial perfusion with 4% PFA. If blood vessels required labelling in any of the three organisms, we injected 10 ml of the dye solution transcardially. Afterwards, organs of interest were removed, and a second quality control was performed by checking if the organ of interest has turned white indicating a successful perfusion. The organs were then incubated overnight at 4°C in 4% PFA, followed by washing three times in 1xPBS for 1hour per wash, and storage in 1xPBS at 4°C.

##### Fish

Preparation of perfusion solutions and dye solutions were the same as for rodents.

Perfusion procedure: Animals were anesthetized in a warm 1:500 2-Phenoxyethanol (2-PE) (Sigma-Aldrich, 77699-250ML) solution. Fish were perfused via cannulation of the outflow tract with ∼50µl of dye solution. Fish were then transferred into ice cold 4%PFA for solidification of gelatin and fixation and overnight incubated at 4°C in 4%PFA followed by three times 1xPBS washes for 1h and storage in 1xPBS at 4°C.

#### Tissue-clearing

##### 3DISCO

Preparation of 3DISCO solutions. Dehydrating solutions were prepared by mixing THF (Sigma-Aldrich 186562-2L) and distilled water (dH_2_O) in the following concentrations: 50 vol% THF, 70 vol% THF, and 80 vol% THF.

The clearing comprised serial incubations of the fixed samples in 10ml of 50 vol% THF, 70 vol% THF (day 1), 80 vol% THF, 100 vol% THF (day 2), 100 vol% THF, and 100 vol% THF (day 3) at 4°C to dehydrate and delipidate the tissue, followed by final RI matching in dibenzyl ether (DBE) (Sigma-Aldrich, 108014-1KG) at 4°C for at least 2 hours until samples became transparent (day 4).

##### FDISCO

Preparation of FDISCO solutions. Dehydrating solutions were prepared by mixing THF (Sigma-Aldrich 186562-2L) and dH_2_O in the following concentrations: 50 vol% THF, 70 vol% THF, and 80 vol% THF.

The clearing comprised serial incubations of the fixed samples in 10 ml of 50 vol% THF, 70 vol% THF (day 1), 80 vol% THF, 100 vol% THF (day 2), 100 vol% THF, and 100 vol% THF (day 3) at 4°C to dehydrate and delipidate the tissue. For each step, THF was adjusted with triethanolamine (Sigma-Aldrich, 90279-100ML) to pH9. The final RI was matched in DBE (Sigma-Aldrich, 108014-1KG) at 4°C for at least 2 hours until samples became transparent (day 4).

##### sDISCO

Preparation of sDISCO solutions. 1L THF purification from peroxides: We filled 4/5 of a chromatography column (Synthware™, C184464C) with aluminum oxide (AL_2_O_3_) (Camag, 5016-A-I). A dropping funnel (Synthware™, F62241L) was placed above the chromatography column, at the bottom of which a collection flask (Synthware™, F41611L) with 2g of butylhydroxytoluene (BHT) was attached. The THF was added to the funnel for purification. For every 1L of THF that is used, ∼ 650-700ml of it will be purified and the rest will be absorbed by the Al_2_O_3_. With this set-up, purified THF contains at least 0.3% BHT, which inhibits peroxide formation during and after the purification process. After purification and final stabilization with BHT, we added 30g Molecular sieves (EMD Millipore, 1.05704.1000) to remove possible water residues, and stored the solution at 4°C.

Dehydrating solutions were prepared by mixing THF and 1xPBS pH9 in the following concentrations: 30 vol% THF, 50 vol% THF, and 70 vol% THF. Further, THF was mixed with dH_2_O in the following concentrations: 80 vol% THF, 90 vol% THF, and 96 vol% THF.

1L DBE purification and stabilization with PG: We placed a filter plate with a pore size of 10-15µm (Synthware™, F601000M) in a filter funnel. Next, we filled 2/3 of the filter funnel with Al_2_O_3_. We then inserted a rubber connector between a collection flask (GrowingLabs, GG5340-1000) and the filter funnel and applied a vacuum to purify DBE. For every liter of DBE, ∼200ml of DBE is absorbed. After purification, we placed the DBE in a dark new flask and added 0.3% PG (Millipore Sigma 02370-100G) to inhibit aldehyde and peroxide formation. We stirred the solution for about one hour to dissolve the PG, and stored the solution at room temperature (RT) or at 4°C.

The clearing comprised serial incubations of the fixed samples in 10ml of 30 vol% THF, 50 vol% THF (day 1), 70 vol% THF, 80 vol% THF (day 2), 90 vol% THF, 96 vol% THF (day 3), 100 vol% THF, and 100 vol% THF (day 4) at 4°C to dehydrate and delipidate the tissue. For the last two dehydration steps, the THF was adjusted with triethanolamine to pH9. The final RI was matched in sDBE at 4°C for at least 2 h until samples became transparent (day 5).

##### uDISCO

Preparation of uDISCO solutions. Dehydrating solutions were prepared by mixing tB (Sigma-Aldrich, 24126-1L) and dH_2_O in the following concentrations: 30 vol% tB, 50 vol% tB, 70 vol% tB, 80 vol% tB, 90 vol% tB, and 96 vol% tB. Delipidation solution dichloromethane (DCM) (Sigma-Aldrich, 34856-100ML) was used as a pure solution. BABB-D4 was prepared by mixing benzyl benzoate (Millipore Sigma, B6630-1L) and benzyl alcohol (Millipore Sigma, 402834-1L) (BABB) in a 2:1 ratio with diphenyl ether (DPE) (Sigma-Aldrich, 67334-25ML) at a ratio of 4:1, and adding 0.4% vol DL-alpha-tocopherol (vitamin E) (Sigma-Aldrich, T3376-25G).

The clearing comprised serial incubations of the fixed samples in 10ml of 30 vol% tB, 50 vol% tB (day 1), 70 vol% tB, 80 vol% tB (day 2), 90 vol% tB, and 100% tB (day 3) at 34-35°C to dehydrate and delipidate the tissue, followed by immersion in DCM for 45-60min at RT for stronger lipid removal and final RI matching in BABB-D4 at RT for at least 2h until samples became transparent (day 4).

##### PEGASOS

Decolorization solution: Quadrol (Millipore Sigma, 122262-1L) and ammonium (Merck, 1054321011) were combined in a single aqueous solution, with final concentrations of 25% (v/v) Quadrol and 5% (v/v) ammonium in H O.

Gradient tB dehydration and delipidating solution: Pure tB (Sigma-Aldrich, 24126-1L) was diluted with dH_2_O to prepare gradient delipidating solutions: 30% v/v, 50% v/v, and 70% v/v. Quadrol (Sigma-Aldrich 122262) was then added with 3% w/v final concentration to adjust the pH to above 9.5. tB-PEG dehydration solution was composed of 70% v/v tB, 27% v/v PEG methacrylate Mn500 (PEG-MA500) (Sigma-Aldrich 409529), and 3% w/v Quadrol (Sigma-Aldrich 122262).

Preparation of polyethylene glycol variants (PEG-X) containing RI media: Three types of polyethylene glycol-methacrylate (PEG-MA) that differ in molecular weight [i.e. PEG-MA200 (Polyscience, 16712-100), PEG-MA440 (Polyscience, 24890-100), and PEG-MA500 (Sigma-Aldrich, 409529-100ML)] were mixed with benzyl benzoate (BB) (Millipore Sigma, B6630-1L or TCI-Chemicals, B0064-500G). PEG-MA was mixed in a 1:9 ratio with BB. The RI of the PEG-MA-BB solutions was adjusted with an AR4 Analog Abbe refractometer (Krüss, K-AR4) to a final RI of 1.543 by adding BB (to increase RI) or PEG-MA (to decrease RI). Depending on the vendor, the BB arrived as a yellow (Sigma Millipore) or colorless (TCI-Chemicals) liquid, which also influenced the final RI medium color.

Clearing comprised two days of incubating the fixed samples in 20ml decolorization solution, followed by serial incubation in 10ml of 30 vol% tB, 50 vol% tB (day 3), 70 vol% tB, 75 vol% tB+ PEG-MA500 (day 4), and 75 vol% tB+ PEG-MA500 (day 5) at 34-35°C to dehydrate and delipidate the tissue, followed by immersion in the final 3%Quadrol:75%BB:22%PEG-MA500 RI-matching solution (day 6) for at least 2 h until samples became transparent. The RI matching solution (3%Quadrol:75%BB:22%PEG-MA500) was changed one more time before imaging (day 7).

##### Testing of Dehydration Agents and RI for Morphological and Clearing Properties of Mouse Brains

Preparation of solutions. Dehydrating solutions were prepared by mixing either methanol (MeOH), ethanol (EtOH), tB, 1-propanol, or THF and dH_2_O in the following concentrations: 50 vol%, 70 vol%, 80 vol%, and 90 vol%.

The clearing comprised serial incubation of the fixed samples in 10 ml of 50 vol%, 70 vol% (day 1), 80 vol%, 90 vol% (day 2), and 2x100 vol% (day 3) to dehydrate and delipidate the tissue. This was followed by serial incubation of samples in tB and 1-propanol at 37°C, MeOH, EtOH, and THF at 4°C. For final RI matching, samples were immersed in either DBE, BABB, or Eci (Sigma-Aldrich, 108014-1KG) at RT for at least 2 h until samples became transparent (day 4).

##### New Tissue-clearing Method for XFP Preservation (XFP-DISCO)

Preparation of solutions for rodent and brain organoids. Solution-1 consists of 8 wt % THEED (Sigma-Aldrich, 87600-100ML), 5 wt % Triton^®^ X 100 (Roth, 3051.2), and 25 wt % urea (Roth, X999.2) in dH_2_O. Dehydrating solutions were prepared by mixing tB and dH_2_O in the following concentrations: 50 vol% tB, 70 vol% tB, 80 vol% tB, and 90 vol% tB+10% polyethylene glycol-methyl ether methacrylate 500 (PEG-MEMA500) (Sigma-Aldrich, 447943-500ML). For each step, pH was adjusted by adding ∼ 3% w/v Quadrol. Dehydration-based RI medium: PEG-MEMA500 was mixed in a 1:9 ratio with sDBE (see preparation in sDISCO protocol) and the final RI was also adjusted to 1.543 by adding sDBE (to increase RI) or PEG-MEMA500 (to decrease RI) using a refractometer. Water-based RI medium (Solution-2): 61% (w/w) meglumine diatrizoate (MD) (Millipore Sigma, M5266-100G) was mixed with dH_2_O under gentle stirring and heating. The final RI was adjusted to 1.45 by adding MD (to increase RI) or dH_2_O (to decrease RI).

The clearing of rodent tissue comprised one-day incubation of the fixed samples in 20ml Solution-1 (day 1), followed by 3x washing in 1xPBS for 1hour each and serial incubations in 10 ml of 50 vol% tB, 70 vol% tB (day 2), 80 vol% tB (day 3), and 90 vol% tB+ 10% PEG-MEMA500 (day 4) at 37°C to dehydrate and delipidate the tissue. The RI was matched with sDBE-PEG-MEMA500 solution for at least 2 h until samples became transparent (day 5).

Optional size modulation: If the final sample size is increased or decreased following completion of the above protocol, alterations to the dehydration procedure should be carried out on day 3. To decrease the sample size: following the day-3 incubation (80 vol% tB), incubate the sample overnight in 90 vol% tB. To increase the sample size: replace the day-3 solution (80 vol% tB) with 90 vol% tB+ 10% PEG-MEMA500. After taking these steps to either decrease or increase the sample size, the remaining steps are unchanged. It must be noted that cortical deformation in brain tissue can occur during dehydration, due to the large percent change in the dehydration solution (70 %tB to 100% tB).

Optional rehydration for water-based clearing: rehydrate the sample via serial incubation in 10ml of 100 vol% tB, 80 vol% tB (day 1), 50 vol% tB, and dH_2_O (day 2) at 35°C and finally RI-match the sample either in 1) Solution-2 in a sealed cuvette, for smaller volumes (<=20ml), or 2), EasyIndex (RI 1.52) (LifeCanvas, EI-500-1.52) in an open cuvette or open chamber, for larger volumes. For imaging, EasyIndex cleared brain were embedded in agarose and immersed in Cargille immersion oil (RI 1.52) to prevent evaporation of EasyIndex and maintain stable RI conditions. Because the oil–sample interface can generate a strong fluorescence boundary, embedding the brain in agarose confined this interface to the oil–agar boundary rather than the tissue itself, thereby enabling artifact-free light-sheet imaging.

For tissue-clearing of brain organoids, we omitted the Solution-1 step and instead started directly with serial incubation in 5ml of 50 vol% tB, 70 vol% tB, 80 vol% tB (day1), and 90 vol% tB+ 10% PEG-MEMA500 (day 2) at 37°C to dehydrate and delipidate the tissue. The final RI was matched with sDBE-PEG-MEMA500 solution at 23-26°C for at least 2 h until samples became transparent (day 3).

##### Tissue-clearing for adult fish

THF and sDBE were purified and stabilized like in the sDISCO protocol. Dehydrating solutions were prepared by mixing tB or THF and dH_2_O in the following concentrations: 50 vol% tB, 70 vol% THF, 80 vol% THF, 90 vol% and 90 vol% THF+10% PEG-MEMA500. sDBE-PEG-MEMA500 RI matching solution was prepared like in the XFP-DISCO protocol.

Fixed and dextran labelled samples were incubated in chilled acetone (-20°C) overnight (day1). The dehydration and dilapidation consisted of serial incubations of the samples in 20ml of 50 vol% tB at 37°C (day 2), 70 vol% THF, 80 vol% THF (day 3), 90 vol% THF, 100 vol% THF (day 4), 90 vol% THF+10% PEG-MEMA500 (day5) at 4°C followed by final RI matching sDBE-PEG-MEMA500 at 23-26°C for at least 2h until samples became transparent (Day 6).

#### Measuring the Effect of Color on Absorption

Brightfield images were taken, under standardized conditions, of agar samples with various concentrations and thicknesses in combination with Rosco filters or mouse brains tissue-cleared under various conditions on the Zeiss AxioZoom stereoscope. Intensity histograms were exported from Zeiss Zen software and converted into heatmaps via GraphPad Prism v10 (Figure 3F and Figures S2J,L). Briefly, intensity in histogram (middle) is derived from 14-bit images and plotted on the scale of 1-6,000. For the heatmaps (right), 14-bit images were converted into 8-bit images and RGB intensities are plotted on the scale of 1-91 (Figure 3E).

#### Measuring Autofluorescence and Fluorescence Signal

Fluorescent images were taken, under standardized conditions, on the above-mentioned stereoscope, of tissue-cleared mouse brains under various conditions. Fluorescent signal and/or autofluorescence were measured by one or more of the following approaches: histogram, arithmetic-mean, maximum-intensity, and heat-map comparison.

Arithmetic mean and maximum intensity comparison: Autofluorescence and signal arithmetic mean and maximum intensity were measured for each step of various clearing protocols and exported from Zeiss Zen software, and the autofluorescence and fluorescence fold change were shown by using GraphPad Prism v10 (Figures 5D,E, Figures S1B, C, E, F, M, N, P, Q, S, T).

Heat map comparison: Autofluorescence intensity histograms were exported from Zeiss Zen software and converted into heatmaps via GraphPad Prism v10 (Figure S2L).

#### Measuring Fluorescence Signal Stability in sDBE

sDISCO tissue-cleared samples were RI-matched with sDBE and imaged with an OMA-LSFM microscope one day after tissue-clearing and 6 months (EYFP) after tissue-clearing. Image stacks were imported into AMIRA, and histograms of fluorescence intensity were generated and overlayed (Figure S1I).

#### Imaging Volumetric Changes with 3D Scanner

Samples were imaged with a 3D scanner with a 0.05-mm accuracy and 0.17-mm resolution (2023 EinScan SP v2 Desktop 3D Scanner) to determine sample volume. Samples were mounted before and after full dehydration on a needle and placed on a turning stage in the 3D scanner. The generated point clouds were exported into AMIRA software, which 3D-visualized the data and generated volumetric values (Fig. 4C, D).

#### Virus Injections

##### Triple Virus Injections

Adeno-associated viral plasmids were obtained from Addgene (pAAV-hSyn1-mTurquoise2, #99125; pAAV-hSyn1-mRuby2, #99126; pAAV-hSyn1-mNeonGreen, #99135) and packaged in-house using a previously published protocol^54^. A total of 500nl of AAV mixture (1:1:1 ratio) was injected in 4-10-month-old C57BL/6J mice at 1.0mm AP, 1.0mm ML, and 0.75mm DV using a microinjector at 10nl/s. Animals were euthanized 4 weeks post-injection and perfused transcardially with 4% PFA for histology or tissue-clearing.

##### Virus Injections in Rats

Adult Long-Evans rats underwent unilateral injections into the ventral posteromedial nucleus (VPm). Coordinates relative to bregma were −3.0mm AP, +2.6mm ML, and −6.0mm DV. Isoflurane anesthesia was induced at 5% and maintained at 2-3%. Following a small craniotomy, a glass pipette was lowered at 10μm/s to the target site.

A total of 200nl of rAAV2/5-hSyn-hChR2(H134R)-EYFP (1.0 × 10¹³ vg/ml) was injected at 0.33nl/s. After a 5-min dwell time, the pipette was retracted 0.5mm, and a second 200-nl injection was performed. The pipette was then slowly withdrawn, and the craniotomy sealed with Kwik-Sil (World Precision Instruments).

##### Virus Injection in *Crh*-Ires^Cre^ Mice

Stereotaxic injections were performed as previously described^55,56^. All surgical procedures were conducted under deep isoflurane anesthesia (5% induction, 1.5–2% maintenance at a flow rate of 1L/min). Mice were secured in a stereotaxic frame (RWD Life Science), and body temperature was maintained using a heating pad throughout the procedure.

Viral particles were delivered using a stereotaxic injector (Stoelting or RWD Life Science) equipped with a pulled glass micropipette (Drummond Scientific). Injections were performed at a controlled rate of 50nl/min. Following infusion, the micropipette was left in place for an additional 5 min before slow withdrawal to minimize backflow and ensure optimal diffusion. Animals were allowed to recover for 21 days to permit sufficient viral expression prior to histological analysis of local and long-range projections.

##### Anterograde Tracing of CRH Projections

To map corticotropin-releasing hormone (CRH) neuronal innervation of the periventricular hypothalamic nucleus, *Crh*-Ires^Cre^ mice (gift from Dr. J.S. Bains, University of Calgary) were used. On postnatal day 21 (P21), mice received a unilateral stereotaxic injection of the Cre-dependent anterograde tracer AAV-hSyn-DIO-EGFP (40nl; Addgene #50457-AAV8) into the paraventricular nucleus of the hypothalamus (PVN). Stereotaxic coordinates relative to bregma were −0.7mm AP, −4.8mm DV, and ±0.2mm ML. Three weeks after injection, animals were deeply anesthetized and transcardially perfused. Brains were collected and processed for light-sheet microscopy to visualize EGFP-labeled projections.

##### Virus Injections in *Chx10*^Cre^ Mice

Mice underwent unilateral viral injections into the left gigantocellular nucleus (Gi) using a Neurostar stereotaxic platform (Tübingen, Germany). Surgeries were conducted under inhaled isoflurane anesthesia (4% induction, ∼2% maintenance), with anesthesia depth monitored via paw pinch. Viral suspensions were mixed with 0.1% fast green dye to visualize delivery during infusion, and injections were performed using pulled glass pipettes. Each injection consisted of 350nl of either AAV5-hSyn1-DIO-hM3D(Gq)-mCherry-WPRE (4 × 10¹² vg/ml, VVF Zurich, lot v89) or AAV5-hSyn1-DIO-hM3D(Gq)-mCitrine-WPRE (2 × 10¹² vg/ml, VVF Zurich, lot 91). Pipettes were left in place for 5 min post-infusion before withdrawal. Target coordinates relative to bregma were −6.0mm AP, −0.8mm ML, and −5.5mm DV^57^. Behavioral testing was performed 3-6 weeks post-surgery. All stereotaxic injections were conducted using standardized coordinates and volumes across replicates to ensure reproducibility.

#### Behavioral Assays for *Chx10* Mice

For open-field analysis, mice were placed in a square 50 × 50 cm chamber under infrared illumination. Video was acquired at 25 frames/s and analyzed with EthoVision XT (Noldus). Tracking in EthoVision XT used head, center, and tail points. The software provided measures of complete 360° body rotations based on changes in the tail-center axis. Rotations were categorized as ipsilateral or contralateral relative to the injected side, and a hysteresis threshold of ±50° was applied within EthoVision to stabilize direction assignment. Each session lasted 60 min, with the initial half-hour for acclimation and the latter half for analysis. For hM3Dq-DREADD experiments, clozapine-N-oxide (CNO; Tocris 4936) was dissolved in saline on the day of testing and delivered intraperitoneally at 0.5mg/kg. Brains from these mice were cleared using our optimized protocol and imaged at 26°C using 12X magnification.

#### Stereomicroscopy

Agar gels and uncleared and tissue-cleared samples, fully immersed in RI medium, were imaged either in brightfield, darkfield, or epifluorescence, all on a stereomicroscope (Zeiss, AxioZoom V16) using a 1xPlanApo Z (NA = 0.25, WD = 60mm) and a Zeiss Axiocam 506 color camera (2752x2208, Pixel size 4.54µm X 4.54µm) with a RGB Bayer color filter mask and a spectral sensitivity ranging from 400nm − 720nm. For some samples, a temperature-controlled imaging chamber was used in combination with the stereomicroscope.

#### Light-sheet Imaging

Light-sheet imaging described in this manuscript was performed using either 1) an Optimized Meso-Aspheric-based light-sheet microscope (OMA-LSFM), 2) a conventional mesoSPIM design, or 3) our mesoSPIM-ultra design. The design of the conventional mesoSPIM is described previously^35^ and is available on https://www.mesoSPIM.org.

##### Light-sheet Imaging with an optimized Meso-aspheric-based light sheet microscope (OMA-LSFM)

The OMA-LSFM microscope used in this work is described partly previously^38^. Briefly, the microscope comprises a dual-sided excitation path using a fiber-coupled multiline laser combiner (488, 532, 594, 647nm, Omicron Light Hub). Either 100% one-sided illumination or a 50% beam splitter is used to divide the incident beam into two identical beams. The beams are guided towards two light-sheet generator units. The details of the light-sheet generators units are (1) aspheric lens (f = 20mm, Linus, Germany), (2) Powell Lens (10° Fan-angle, Edmund Optics, Germany), (3) aspheric lens (f = 20mm, Linus, Germany), (4) Bullseye filter with elliptical aperture (Reynard Cooperation, USA), (5) acylindrical lens (f = 80mm, Linus, Germany), and (6) acylindrical lens (f = 80mm, Linus, Germany). The description of the light-sheet generator is given in the patent DE 102010046133B4 and in Saghafi et al. 2018^58^. The light-sheet generators are placed on a computer-controlled linear stage (LS-65, PI-Micos, Germany) that can be moved along the beam-propagation axis, generating a light sheet with an optimal line of focus propagating through the sample. A computer-controlled elevation stage with an adjustable precision less than 100nm (Es-100, PI-Micos GmbH, Germany), and a manually adjustable xy-cross table for horizontal adjustment, are used to scan the sample vertically.

The light-detection component of the OMA-LSFM includes a customized microscope with modified Evident-Olympus objectives for RI mismatch 2x (XLFluor2x/340, NA = 0.14, WD = 21mm), 4x (XLFluor4x/340, NA = 0.28, WD = 29.5mm), a 16x Leica objective (HC FLUOTAR L 16x NA, 0.6 IMM CORR, WD = 2.5mm), a scientific CMOS (sCMOS) camera (Andor Neo: 2560x2160, Pixel size 6.5μm), and a filter set of a) 460 LP ET long pass-filter F76-460, b) 525/20 ET Bandpass F47-528, and c) 605/70 ET Bandpass F47-605 (all AHF, Germany). The objectives and the samples are immersed from above in a 50×50×50mm³ immersion cuvette.

##### Light-sheet Imaging with Our mesoSPIM-ultra System

The excitation path in our mesoSPIM-ultra is identical to that in the original published mesoSPIM-V5^35^. Briefly, the microscope comprises a dual-sided excitation path using a fiber-coupled multiline laser combiner (LuxX 405nm, 120mW; LuxX 488nm, 150mW; DPSS 561nm, 150mW; DPSS 594nm, 100mW; LuxX 647nm, 140mW in Omicron Light Hub). Galvo scanners (Thorlabs GVS011) are used for light-sheet generation and reduction of streaking artifacts caused by absorption of the light sheet. In addition, the beam waist is scanned using electrically tunable lenses (ETL, Optotune EL-16-40-5D-TC-L) synchronized with the rolling shutter of the sCMOS camera. This axially scanned light-sheet mode (ASLM) leads to a uniform axial resolution across the field-of-view of 5-10 µm (depending on objective and wavelength). The detection path comprises three Miltenyi Biotec objectives (1.1x NA = 0.1, WD = 16mm; 4x NA = 0.35, WD = 15mm; 12x NA = 0.53, WD = 10mm) in a horizontal objective turret actuated by a PI MICOS L-611 rotation stage. The light path is folded using two elliptic 2” mirrors (Thorlabs BBE2-E02) and uses an Olympus SWTLU-C wide-field tube lens (Evident), a ZWO 7 x 2” filter wheel with a filter set of ET480/30m, ET525/30m, ET540/30m, ET605/40m, ET625/40m, and ET665lp (all Chroma, USA), and a sCMOS camera (Teledyne Photometrics Kinetix: 3200x3200, pixel size 6.5μm). Sample translation and rotation is accomplished by an XYZ stage set of three PI L-509 stages with a 100-mm travel range in combination with a M-060.DG rotation stage. The samples are suspended from above in a 40×40×40-mm³ or 50×50×50-mm³ temperature-controlled immersion cuvette that can be translated along the detection axis by a PI L-509 stage. Sample, cuvette, and focusing stages are controlled by a PI C-884.6DC controller. The objective turret stage is controlled by a PI C-863.11 controller. All microscope functions are controlled by a version of the mesoSPIM-control software written in Python (https://github.com/ffvoigt/mesoSPIM-control/tree/mdibl-master).

The assembly and alignment of a standard mesoSPIM is extensively documented in the mesoSPIM wiki (https://github.com/mesoSPIM/mesoSPIM-hardware-documentation/wiki), which also contains documentation for the modifications for integrating the Miltenyi Biotec objectives (https://github.com/mesoSPIM/mesoSPIM-hardware-documentation/wiki/mdibl_overview)

#### Image Processing and Visualization of Light-sheet Data

Image processing and 3D reconstruction was performed using a Lenovo ThinkStation P920 workstation equipped with 2 x Intel® Xeon® Platinum 8280 Processor (2.70 GHz, up to 4.00 GHz Max Boost, 28 cores, 38.5MB Cache), 1TB Ram DDR4 2933Mhz MHz, 16 DIMMs, and 2 x NVIDIA® RTX™ A6000 with NVLink 48GB GDDR6.

CLAHE and the Fast-Fourier-Transform-based destriping algorithm were performed with Wiener software developed by Klaus Becker^59,60^. Image restoration by deconvolution was performed by using either Wiener software or Huygens Professional 24.10 (Scientific Volume Imaging, Netherlands) and where applicable hot pixel removal, Classic Maximum Likelihood Estimation (CMLE) and SNR estimation.

Image registration and stitching were performed with Imaris Sticher 10.2.0 (Oxford Instruments, UK) or AMIRA 2022.1 (Thermo Fisher Scientific, USA). 3D Visualization was performed either by Imaris 10.2.0, Amira 2022, or syGlass v2.1.0 virtual reality (VR) (syGlass, USA) software.

Counting and tracing of *Chx10*^+^ neurons was performed manually in VR by using MetaQuest 3 VR goggles (Meta, USA) with the counting and tracing tool in the syGlass software (Figures 7E and G-K).

### QUANTIFICATION AND STATISTICAL ANALYSIS

#### Total Transparency and Haze Measurement and Quantification

For each measurement using the hazemeter (HFBTE, Digital Glass hazemeter), we calibrated the instrument before taking the measurement, as follows. Total transmission (TT) and haze (H) were calibrated by placing agar (Figures 1C-M, 3B-D) or tissue-cleared samples (fully immersed), separately, in the same amount of RI medium in separate temperature-controlled cuvettes (Figures 1N-Q, 2B, 4E-G, 5F, S2F-I), and then placing each cuvette, separately, on the hazemeter. Mann-Whitney test was used to evaluate the effects of the condition, i.e., the agar or tissue-cleared samples. Normality of the data was not assumed. Data are presented as box-and-whisker plots (median, quartiles, range). Significance was defined as P < 0.05, with thresholds indicated as not significant (ns) P > 0.05, * P ≤ 0.05, ** P ≤ 0.01, *** P ≤ 0.001, and **** P ≤ 0.0001. GraphPad Prism v10 was used for all analyses.

#### Quantification of Sample Volume Changes with a Pycnometer

The pycnometer (Bigamart, B07QFRKD3K) was filled with dH_2_O, and the total weight of the pycnometer and dH_2_O was measured on a fine scale. The sample was then placed inside the pycnometer and the lid to the pycnometer was closed, resulting in displacement of excess water, via the Archimedes principle, through a fine capillary in the lid. The sample was then removed, and the total weight of the pycnometer and the remaining water was measured again. The difference between the two measurements indicated the volume of the sample. This was performed for each dehydration step of each sample (Fig. 4B). Mann-Whitney test was used to evaluate the effects of the dehydration condition. Normality of the data was not assumed. Data are presented as box-and-whisker plots (median, quartiles, range). Significance was defined as P < 0.05, with thresholds indicated as not significant (ns) P > 0.05, * P ≤ 0.05, ** P ≤ 0.01 and *** P ≤ 0.001. GraphPad Prism v10 was used for all analyses.

#### Statistical Analysis of Turning Behavior in *Chx10* Mice

Group assignments were block-randomized. Sample sizes were consistent with those in prior publication^40^ but not predetermined by power analysis. All data were included, regardless of whether or not mice showed behavioral responses to CNO. Behavioral scoring was automated; thus, while experimenters were aware of group identity, operator bias was minimized. Two-way ANOVAs were used to evaluate effects of condition, with Tukey-Kramer HSD tests for post hoc comparisons. Normality of the data was assumed but not formally tested. Data are presented as box-and-whisker plots (median, quartiles, range) with individual values overlaid. Significance was defined as P < 0.05, with thresholds indicated as P < 0.05, *P < 0.01, and **P < 0.001. GraphPad Prism v10 was used for all analyses.

### KEY RESOURCES TABLE

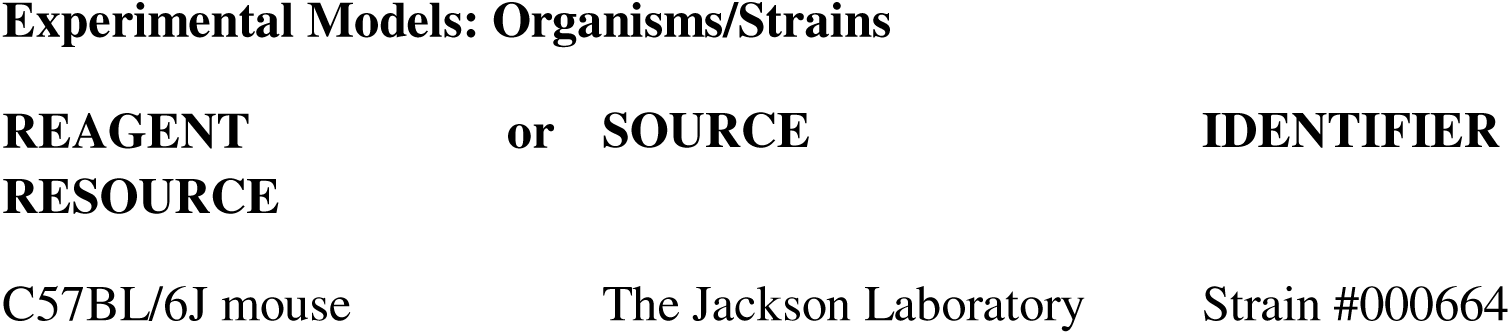
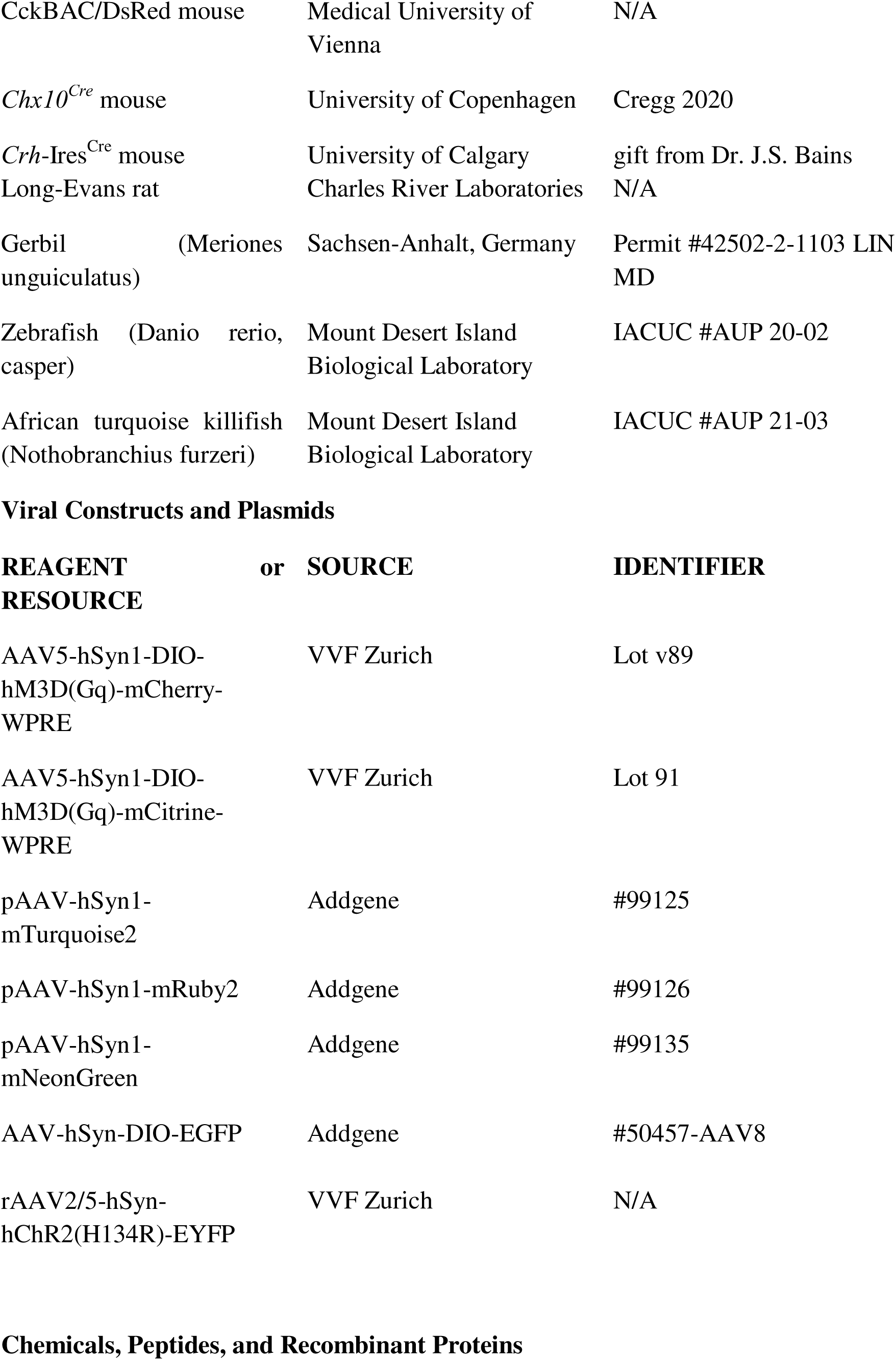
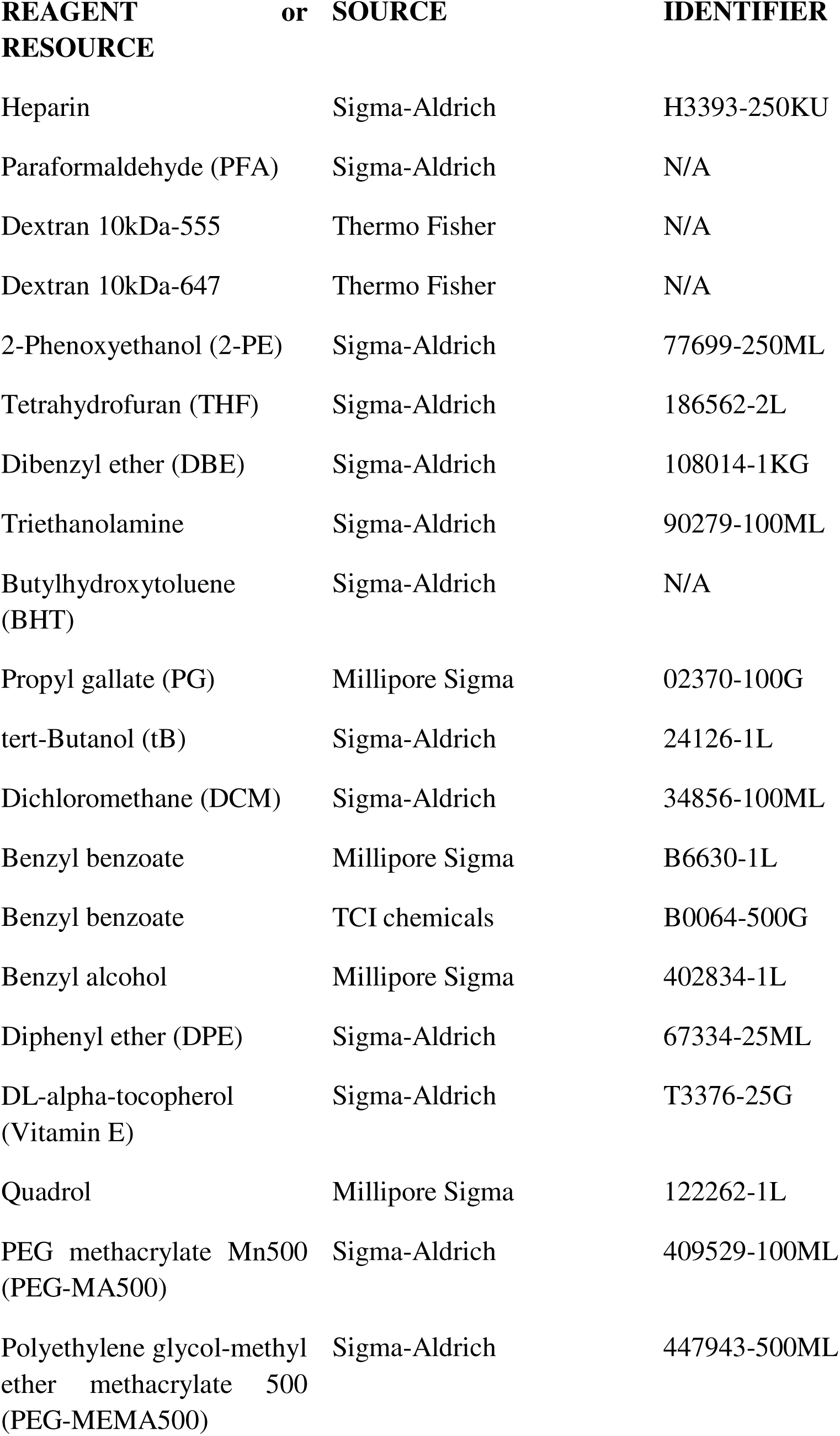
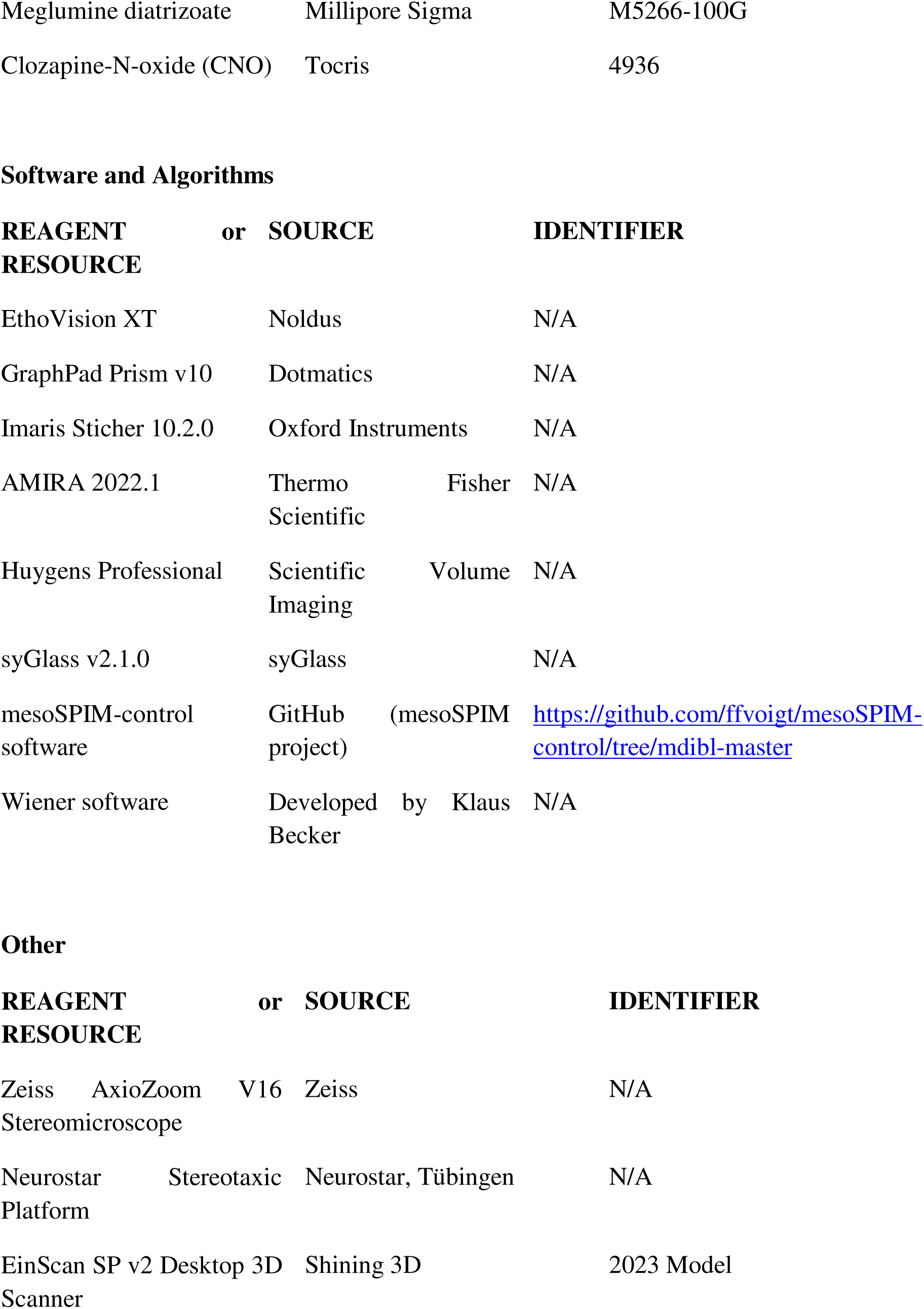
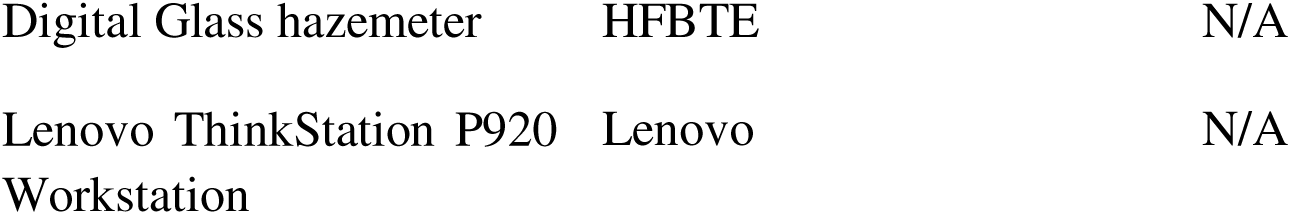

---

**Supplementary Figure 1.**
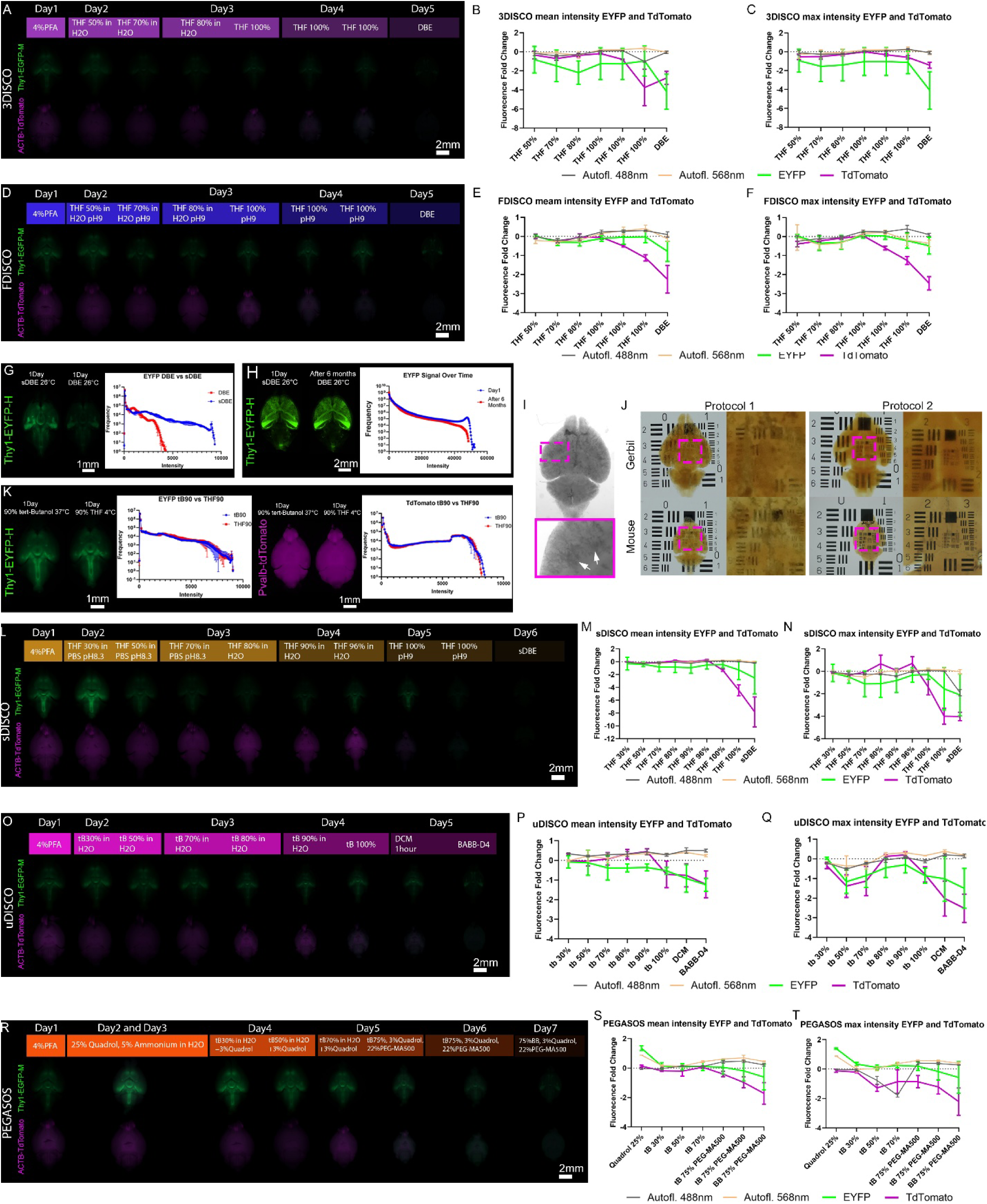
Comparative Study of Mouse-brain Clearing Protocols for XFP Preservation. (A) Overview of the 3DISCO tissue-clearing protocol with representative epifluorescence images showing fluorescent-signal retention at each step. Abbreviations: PFA: paraformaldehyde; THF: tetrahydrofuran; DBE: dibenzyl ether. (B–C) Fold changes in mean (B) and maximum (C) autofluorescence and transgenic fluorescent signal across 3DISCO protocol steps, normalized to PFA-fixed baseline levels (n = 3 brains). (D) Overview of the FDISCO tissue-clearing protocol with representative epifluorescence images showing fluorescent-signal retention at each step. (E–F) Fold changes in mean (E) and maximum (F) autofluorescence and transgenic fluorescent signal across FDISCO protocol steps, normalized to PFA-fixed baseline levels (n = 3 brains). (G) Epifluorescence images showing a decrease in fluorescent signal in tissue-cleared Thy1*–*EYFP–H (green) mouse brains after transfer from sDBE to DBE (one day in each medium), with corresponding signal-intensity histogram. (H) Impact of sDBE on fluorescent-signal retention. Light-sheet images showing fluorescent signals in tissue-cleared Thy1–EYFP–H (green) mouse brain one day after clearing and 6 months post-clearing (EYFP stable up to this point), with corresponding signal-intensity histogram. (I, J) Brightfield images of tissue-cleared brains using stabilized dibenzyl ether (sDBE) for refractive-index (RI) matching. (I) Grayscale image showing precipitation within blood vessels. Insets show higher-magnification views of boxed regions with arrows indication precipitation in blood vessels. (J) Left: Precipitation in blood vessels of rodent brains obstructing tissue transparency when using the sDISCO protocol without tB (Protocol 1: previously published protocol). Right: absence of precipitation after inclusion of a tB modification step in the sDISCO protocol (Protocol 2: modified protocol). Insets show higher-magnification views of boxed regions. (K) Epifluorescence images showing fluorescent-signal changes in Thy1–EYFP–H (green) and Pvalb–tdTomato (magenta) mouse brains after one day in 90% tB followed by one day in 90% THF, with corresponding signal-intensity histograms. (L) Overview of the sDISCO tissue-clearing protocol with representative epifluorescence images showing fluorescent-signal retention at each step. Abbreviation: sDBE: stabilized dibenzyl ether. (M, N) Fold changes in mean (M) and maximum (N) autofluorescence and transgenic fluorescent signal across FDISCO protocol steps, normalized to PFA-fixed baseline levels (n = 3 brains). (O) Overview of the uDISCO tissue-clearing protocol with representative epifluorescence images showing fluorescent-signal retention at each step. Abbreviations: tB: tert-Butanol; DCM: dichloromethane; BABB-D4: benzyl ether/benzyl benzoate with diphenyl ether. (P, Q) Fold changes in mean (P) and maximum (Q) autofluorescence and transgenic fluorescent signal across uDISCO protocol steps, normalized to PFA-fixed baseline levels (n = 3 brains). (R) Overview of the PEGASOS tissue-clearing protocol with representative epifluorescence images showing fluorescent signal retention at each step. Abbreviations: PEG-MA500: polyethylene glycol methacrylate 500; BB: benzyl benzoate. (S, T) Fold changes in mean (S) and maximum (T) autofluorescence and transgenic fluorescent signal across PEGASOS protocol steps, normalized to PFA-fixed baseline levels (n = 3 brains).

**Supplementary Figure 2.**
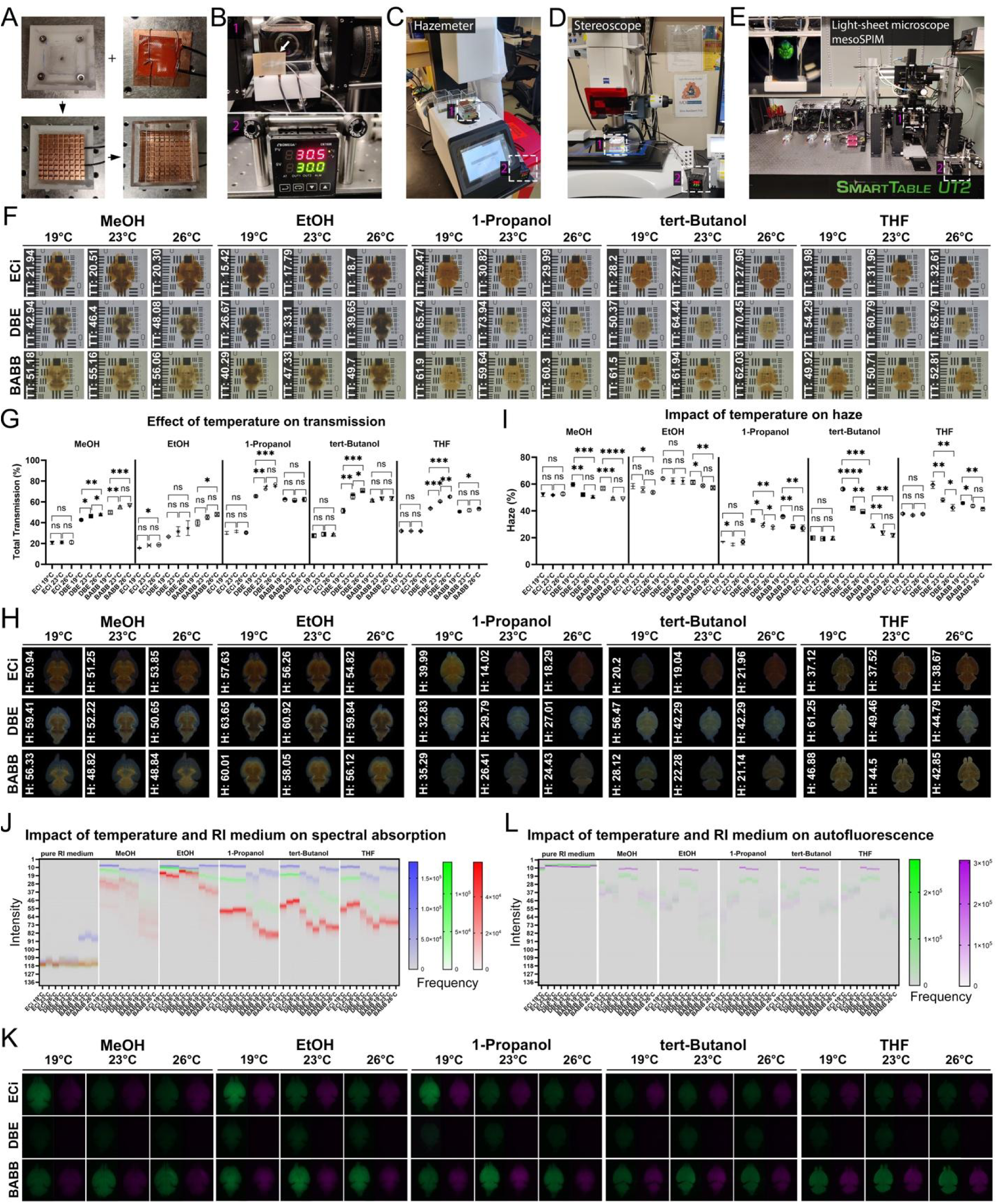
Temperature-dependent Transparency, Spectral Absorption, and Autofluorescence in Mouse Brains Cleared with Different Dehydration and Refractive-Index Agents. (A) Assembly of the heating unit and imaging chamber. From left to right: base; heatsink with mounted heating pad; integration of base, heatsink, and heating pad; glass cuvette positioned on the heatsink. (B) Temperature control components: (1) temperature probe; (2) heating unit. Arrow point at temperature measurement probe. (C–E) Integration of the heated imaging chamber with (C) a hazemeter, (D) a stereoscope, and (E) a light-sheet microscope (F-I) Impact of temperature, dehydration agent, and refractive index (RI)-matching medium on total transmission (TT) and haze (H) using mouse brains. Dehydration agents: methanol (MeOH), ethanol (EtOH), tert-Butanol (tB), 1-Propanol, and tetrahydrofuran (THF); RI-matching media: ethyl cinnamate (ECi), dibenzyl ether (DBE), and benzyl alcohol/benzyl benzoate (BABB). (F, H) Qualitative transparency of mouse brain assayed via brightfield imaging on USAF chart, and TT values measured via hazemeter (F); qualitative assessment of light scattering assayed via darkfield imaging of mouse brain; and H values measured via hazemeter (H). Rows show the same brain imaged at different temperatures. (G) Data are plotted based on TT values from (F). (ns: not significant, P > 0.05, * P ≤ 0.05, ** P ≤ 0.01, *** P ≤ 0.001, Mann-Whitney test). (I) Data are plotted based on H values from (H). (** P ≤ 0.01, *** P ≤ 0.001,**** P ≤ 0.0001, Mann-Whitney test). (J) Heatmap representations of RGB histograms corresponding to the conditions shown in (F). Same sized ROI within the brain were used for measurement. For the heatmap 14-bit images of RGB histograms were converted into 8-bit images (0-255 gray values) and RGB intensities are plotted on the scale up to 136 gray values. (K) Impact of temperature, dehydration agent, and refractive-index (RI)-matching medium on autofluorescence at 488 nm (green) and 568 nm (magenta), using wild-type mouse brains. Dehydration agents: methanol (MeOH), ethanol (EtOH), tert-Butanol (tB), 1-Propanol, and tetrahydrofuran (THF); RI-matching media: ethyl cinnamate (ECi), dibenzyl ether (DBE), and benzyl alcohol/benzyl benzoate (BABB). (L) Heatmap representations of brain-tissue autofluorescence at 488 nm (green) and 568 nm (magenta) based on the data shown in (K), Same sized ROI within the brain were used for measurement. For the heatmap 14 bit images of grayscale histograms were converted into 8bit images (0-255 gray values) and intensities are plotted on the scale up to 136 gray values.

**Supplementary Figure 3.**
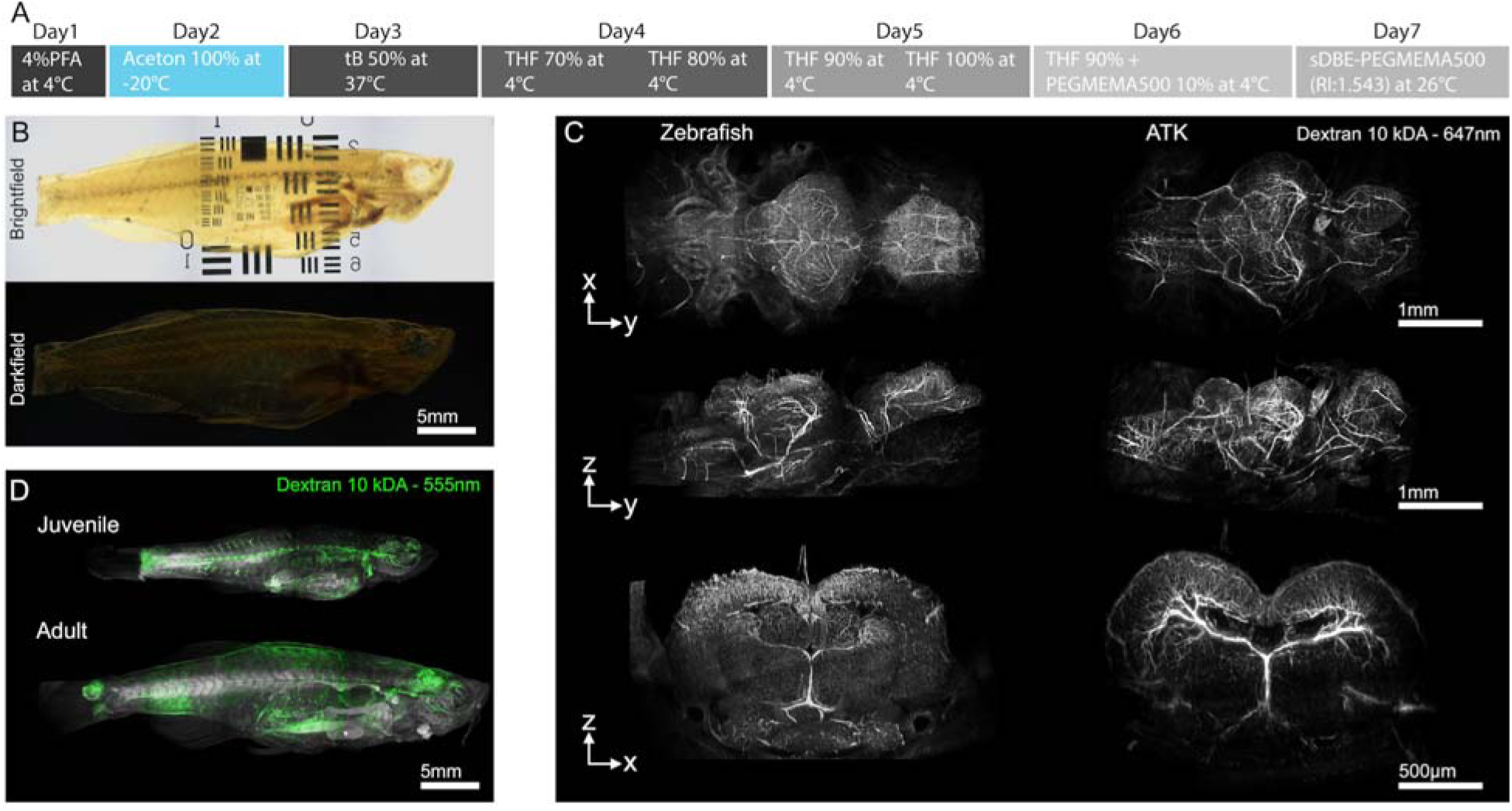
Blood-vessel Visualization in Zebrafish and African Turquoise Killifish. (A) Tissue-clearing protocol for whole adult zebrafish. PFA: paraformaldehyde, tB: tert-Butanol, THF: tetrahydrofuran, sDBE: stabilized dibenzyl ether, PEGMEMA500: polyethylene glycol methyl ether methacrylate 500 (B) Brightfield (top) and darkfield (bottom) images of adult casper zebrafish. (C–E) Light-sheet images of tissue-cleared samples showing vasculature. (C) Brain of juvenile zebrafish (5 months old) and African turquoise killifish (ATK) (5 weeks old) imaged with a 16× (NA-0.6) objective showing vasculature. Image on OMA-LSFM light-sheet microscope. (D) Whole-body imaging of juvenile (5 months old, top) and adult zebrafish (20 months old, bottom) using a 2× NA-0.14 objective. Image on OMA-LSFM light-sheet microscope.

**Supplementary Figure 4.**
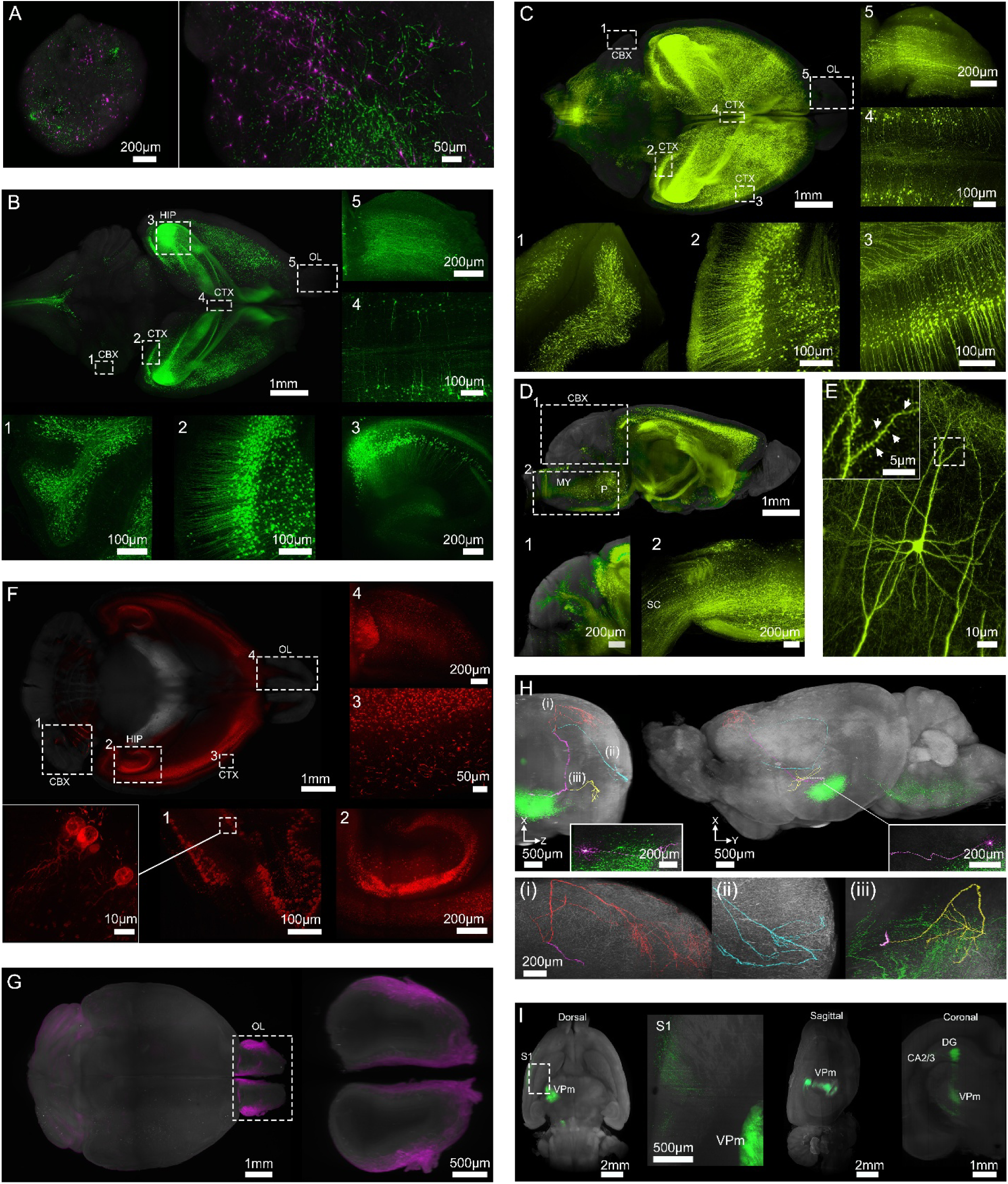
XFP Signal Preservation in Tissue-cleared Samples. (A–I) Light-sheet images of tissue-cleared mouse brains acquired on an OMA-LSFM system. (A) Brain organoid showing CAG:GFP (green) and CAG:tdTomato (magenta) fluorescence (left). High magnification of neuronal rosettes (right) (B) Whole-brain dorsal view of a Thy1–EGFP–M mouse brain. Insets show higher-magnification views of dashed areas highlighting neuronal projections in the cerebellum (1), cortex (2), hippocampus (3), cortical midline between hemispheres (4), and olfactory lobe (5). (C) Whole-brain dorsal view of a Thy1–EYFP–H mouse brain. Insets show higher-magnification views of dashed areas highlighting neuronal projections in the cerebellum (1), cortical areas (2 and 3), cortical midline between hemispheres (4), and olfactory lobe (5). (D) Whole-brain sagittal view of a Thy1–EYFP–H mouse brain. Insets show higher-magnification views of dashed areas highlighting neuronal projections in the cerebellum (1) and spinal cord (2). (E) Cortical neuron in a Thy1–EYFP–H mouse brain, with an inset showing higher-magnification view of the dashed area; arrows indicate dendritic spines. (F) Dorsal view of a Cck^BAC/DsRed^ mouse brain optical section, showing cholecystokinin (Cck)-positive neurons in the cerebellar Purkinje cell layer (1), hippocampal pyramidal layer (2), cortex (3), and olfactory lobe (4). Inset in (3) shows a higher-magnification view of labeled neurons. inset shows a higher-magnification of basket cells in the cerebellum (G) Whole-brain dorsal view of a brain of Pvalb–TdTomato mice; higher-magnification inset of the dashed area shows the olfactory lobes. (H) Expression of AAV-DIO-EGFP tracer injected into the dorsal hypothalamic area of a *CRH*-Ires-Cre mouse brain. Upper panel: intra-hypothalamic arborization of CRH neurons (magenta) and long-range axonal projections to the (i) somatosensory cortex (red), (ii) somatomotor (cyan), (iii) ventrolateral thalamic nucleus (yellow), and trigeminal nucleus in the medulla (green). Insets show higher magnification of neuronal origin. Lower panel: Lateral resolution of color-coded axonal projections. (I) Rat brain injected with 400nl of rAAV2/5-hSyn-hChR2(H134R)-EYFP (titer: 1.0 × 10¹³ vg/ml). *Dorsal view:* Ventral posteromedial nucleus (VPm) of the thalamus projecting to the primary somatosensory (S1) cortex. Inset shows a higher-magnification view of the dashed rectangle. *Sagittal and coronal views:* Expression to the right of the hippocampus reveals dentate gyrus (DG) projections to CA2/3 regions. CTX: Cortex, CBX: Cerebellum, HIP: Hippocampus, OL: Olfactory lobe, P: Pons, MY: Medulla, SC: Spinal Cord

**Supplementary Figure 5.**
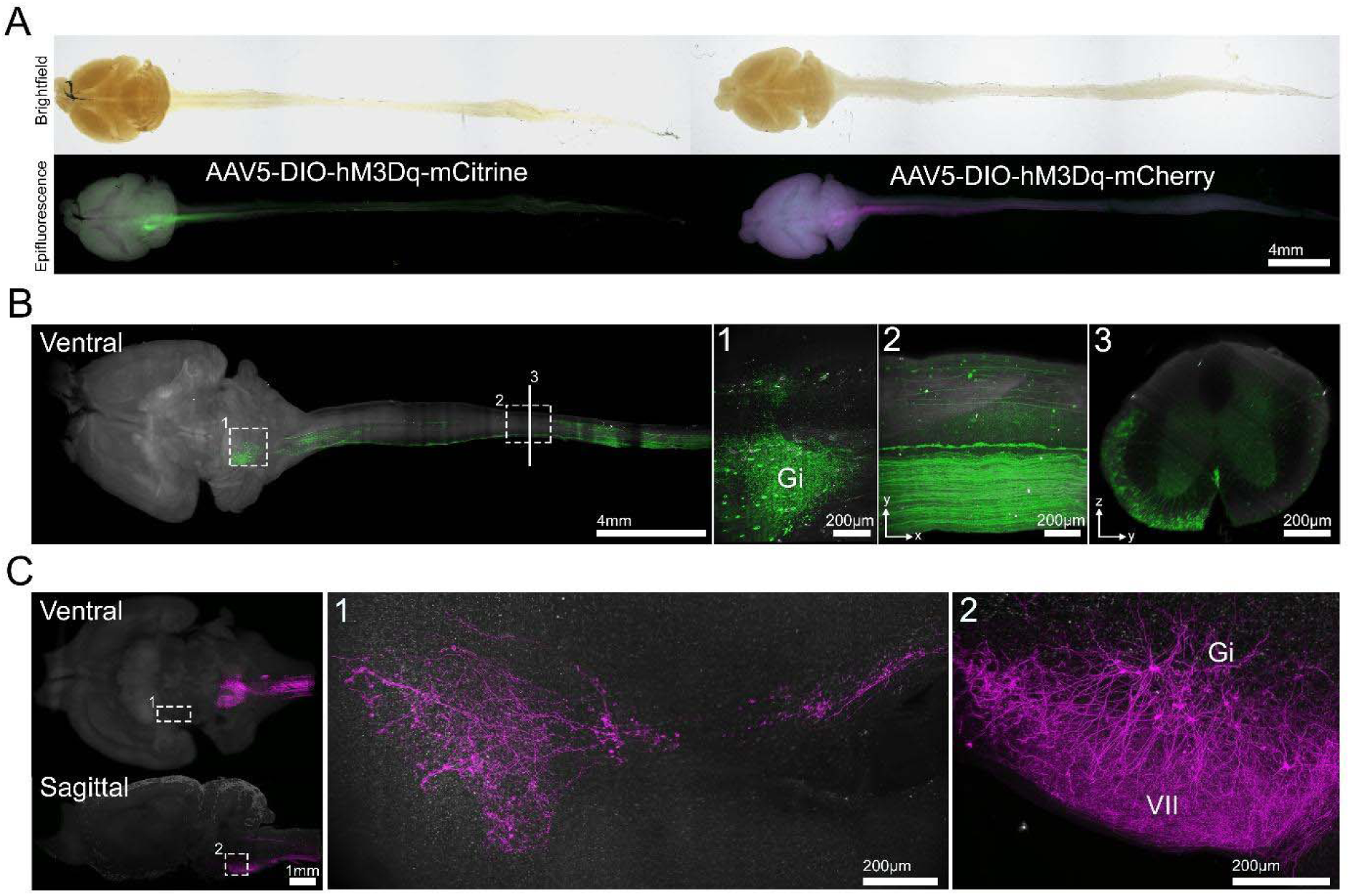
Visualization of the *Chx10* Gi Projectome. (A) Top: Brightfield images of a tissue-cleared *Chx10^Cre^* mouse central nervous system (CNS: brain and spinal cord). Bottom: Corresponding epifluorescence images showing distinct fluorescent protein expression (mCitrine (in green) (left) and mCherry (in magenta) (right) following viral injection. (B) Light-sheet images of a tissue-cleared *Chx10^Cre^* mouse injected with 350 nl AAV5-DIO-hM3Dq-mCitrine (titer: 2.0 x 10^12^ vg/ml). Insets: (1) injection site in Gi; (2) axonal projections within the spinal cord; (3) coronal spinal-cord section showing axon profiles in the ventrolateral funiculus and axonal arborizations in the ventral horn. Gi: Gigantocellular reticular nucleus. (C) Light-sheet images of a tissue-cleared *Chx10^Cre^* mouse CNS-injected with 350 nl AAV5-DIO-hM3Dq-mCherry in the left Gi (titer: 4.0 x 10^12^ vg/ml). Insets: (1) sparse midbrain projections of *Chx10* neurons; (2) injection site in Gi and innervation of VII. VII: facial motor nucleus.

Related to Main Figure 5 - Supplementary Video 1:

Tissue-cleared mouse brain showing intra-hypothalamic arborization of CRH neurons (magenta) and long-range axonal projections to the somatosensory cortex (red), somatomotor (cyan), ventrolateral thalamic nucleus (yellow), and trigeminal nucleus in the medulla (green) using OMA-LSFM with a 4× NA-0.28 and 16x NA-0.6 objective.

Related to Main Figure 6 - Supplementary Video 2:

Whole-body imaging of autofluorescence (grey) and 10 kDA dextran 555 nm (magenta) labelled vasculature in tissue-cleared adult zebrafish (20 months old) using mesoSPIM-ultra with a 4× NA-0.35 objective.

Related to Main Figure 7 - Supplementary Video 3:

*En block* CNS imaging of tissue-cleared mouse brain showing *Chx10^+^* neuronal projections expressing TdTomato using mesoSPIM-ultra with a 12× NA-0.53 objective. Axonal projections are rendered with syGlass VR software.

## REFERENCES

1. Ueda, H.R., Erturk, A., Chung, K., Gradinaru, V., Chedotal, A., Tomancak, P., and Keller, P.J. (2020). Tissue clearing and its applications in neuroscience. Nat Rev Neurosci 21, 61–79. 10.1038/s41583-019-0250-1.

2. Richardson, D.S., and Lichtman, J.W. (2015). Clarifying Tissue Clearing. Cell 162, 246–257. 10.1016/j.cell.2015.06.067.

3. Vieites-Prado, A., and Renier, N. (2021). Tissue clearing and 3D imaging in developmental biology. Development 148. 10.1242/dev.199369.

4. Pende, M., Vadiwala, K., Schmidbaur, H., Stockinger, A.W., Murawala, P., Saghafi, S., Dekens, M.P.S., Becker, K., Revilla, I.D.R., Papadopoulos, S.C., et al. (2020). A versatile depigmentation, clearing, and labeling method for exploring nervous system diversity. Sci Adv 6, eaba0365. 10.1126/sciadv.aba0365.

5. Pende, M., Saghafi, S., Becker, K., Hummel, T., and Dodt, H.U. (2022). FlyClear: A Tissue-Clearing Technique for High-Resolution Microscopy of Drosophila. Methods Mol Biol 2540, 349–359. 10.1007/978-1-0716-2541-5_18.

6. Neckel, P.H., Mattheus, U., Hirt, B., Just, L., and Mack, A.F. (2016). Large-scale tissue clearing (PACT): Technical evaluation and new perspectives in immunofluorescence, histology, and ultrastructure. Sci Rep 6, 34331. 10.1038/srep34331.

7. Tomer, R., Ye, L., Hsueh, B., and Deisseroth, K. (2014). Advanced CLARITY for rapid and high-resolution imaging of intact tissues. Nat Protoc 9, 1682–1697. 10.1038/nprot.2014.123.

8. Ren, Z., Wu, Y., Wang, Z., Hu, Y., Lu, J., Liu, J., Chen, Y., and Yao, M. (2021). CUBIC-plus: An optimized method for rapid tissue clearing and decolorization. Biochem Biophys Res Commun 568, 116–123. 10.1016/j.bbrc.2021.06.075.

9. Pinheiro, T., Mayor, I., Edwards, S., Joven, A., Kantzer, C.G., Kirkham, M., and Simon, A. (2021). CUBIC-f: An optimized clearing method for cell tracing and evaluation of neurite density in the salamander brain. J Neurosci Methods 348, 109002. 10.1016/j.jneumeth.2020.109002.

10. Matsumoto, K., Mitani, T.T., Horiguchi, S.A., Kaneshiro, J., Murakami, T.C., Mano, T., Fujishima, H., Konno, A., Watanabe, T.M., Hirai, H., and Ueda, H.R. (2019). Advanced CUBIC tissue clearing for whole-organ cell profiling. Nat Protoc 14, 3506–3537. 10.1038/s41596-019-0240-9.

11. Ke, M.T., Fujimoto, S., and Imai, T. (2013). SeeDB: a simple and morphology-preserving optical clearing agent for neuronal circuit reconstruction. Nat Neurosci 16, 1154–1161. 10.1038/nn.3447.

12. Hildebrand, S., Schueth, A., Wangenheim, K.V., Mattheyer, C., Pampaloni, F., Bratzke, H., Roebroeck, A.F., and Galuske, R.A.W. (2020). hFRUIT: An optimized agent for optical clearing of DiI-stained adult human brain tissue. Sci Rep 10, 9950. 10.1038/s41598-020-66999-3.

13. Kuwajima, T., Sitko, A.A., Bhansali, P., Jurgens, C., Guido, W., and Mason, C. (2013). ClearT: a detergent- and solvent-free clearing method for neuronal and non-neuronal tissue. Development 140, 1364–1368. 10.1242/dev.091844.

14. Hama, H., Hioki, H., Namiki, K., Hoshida, T., Kurokawa, H., Ishidate, F., Kaneko, T., Akagi, T., Saito, T., Saido, T., and Miyawaki, A. (2015). ScaleS: an optical clearing palette for biological imaging. Nat Neurosci 18, 1518–1529. 10.1038/nn.4107.

15. Li, W., Germain, R.N., and Gerner, M.Y. (2019). High-dimensional cell-level analysis of tissues with Ce3D multiplex volume imaging. Nat Protoc 14, 1708–1733. 10.1038/s41596-019-0156-4.

16. Gao, Y., Xin, F., Wang, T., Shao, C., Hu, Y., Chen, Z., Wang, Y., Xie, F., Li, T., Li, S., et al. (2025). VIVIT: Resolving trans-scale volumetric biological architectures via ionic glassy tissue. Cell 188, 6079–6095 e6020. 10.1016/j.cell.2025.07.023.

17. Louvel, V., Haase, R., Mercey, O., Laporte, M.H., Eloy, T., Baudrier, E., Fortun, D., Soldati-Favre, D., Hamel, V., and Guichard, P. (2023). iU-ExM: nanoscopy of organelles and tissues with iterative ultrastructure expansion microscopy. Nat Commun 14, 7893. 10.1038/s41467-023-43582-8.

18. Gambarotto, D., Zwettler, F.U., Le Guennec, M., Schmidt-Cernohorska, M., Fortun, D., Borgers, S., Heine, J., Schloetel, J.G., Reuss, M., Unser, M., et al. (2019). Imaging cellular ultrastructures using expansion microscopy (U-ExM). Nat Methods 16, 71–74. 10.1038/s41592-018-0238-1.

19. Masselink, W., Reumann, D., Murawala, P., Pasierbek, P., Taniguchi, Y., Bonnay, F., Meixner, K., Knoblich, J.A., and Tanaka, E.M. (2019). Broad applicability of a streamlined ethyl cinnamate-based clearing procedure. Development 146. /10.1242/dev.166884.

20. Becker, K., Jahrling, N., Saghafi, S., Weiler, R., and Dodt, H.U. (2012). Chemical clearing and dehydration of GFP expressing mouse brains. PLoS One 7, e33916. 10.1371/journal.pone.0033916.

21. Erturk, A., Becker, K., Jahrling, N., Mauch, C.P., Hojer, C.D., Egen, J.G., Hellal, F., Bradke, F., Sheng, M., and Dodt, H.U. (2012). Three-dimensional imaging of solvent-cleared organs using 3DISCO. Nat Protoc 7, 1983–1995. 10.1038/nprot.2012.119.

22. Renier, N., Wu, Z., Simon, D.J., Yang, J., Ariel, P., and Tessier-Lavigne, M. (2014). iDISCO: a simple, rapid method to immunolabel large tissue samples for volume imaging. Cell 159, 896–910. 10.1016/j.cell.2014.10.010.

23. Pan, C., Cai, R., Quacquarelli, F.P., Ghasemigharagoz, A., Lourbopoulos, A., Matryba, P., Plesnila, N., Dichgans, M., Hellal, F., and Erturk, A. (2016). Shrinkage-mediated imaging of entire organs and organisms using uDISCO. Nat Methods 13, 859–867. 10.1038/nmeth.3964.

24. Qi, Y., Yu, T., Xu, J., Wan, P., Ma, Y., Zhu, J., Li, Y., Gong, H., Luo, Q., and Zhu, D. (2019). FDISCO: Advanced solvent-based clearing method for imaging whole organs. Sci Adv 5, eaau8355. 10.1126/sciadv.aau8355.

25. Cai, R., Kolabas, Z.I., Pan, C., Mai, H., Zhao, S., Kaltenecker, D., Voigt, F.F., Molbay, M., Ohn, T.L., Vincke, C., et al. (2023). Whole-mouse clearing and imaging at the cellular level with vDISCO. Nat Protoc 18, 1197–1242. 10.1038/s41596-022-00788-2.

26. Hahn, C., Becker, K., Saghafi, S., Pende, M., Avdibasic, A., Foroughipour, M., Heinz, D.E., Wotjak, C.T., and Dodt, H.U. (2019). High-resolution imaging of fluorescent whole mouse brains using stabilised organic media (sDISCO). J Biophotonics 12, e201800368. 10.1002/jbio.201800368.

27. Zhao, S., Todorov, M.I., Cai, R., Maskari, R.A., Steinke, H., Kemter, E., Mai, H., Rong, Z., Warmer, M., Stanic, K., et al. (2020). Cellular and Molecular Probing of Intact Human Organs. Cell 180, 796–812 e719. 10.1016/j.cell.2020.01.030.

28. Jing, D., Zhang, S., Luo, W., Gao, X., Men, Y., Ma, C., Liu, X., Yi, Y., Bugde, A., Zhou, B.O., et al. (2018). Tissue clearing of both hard and soft tissue organs with the PEGASOS method. Cell Res 28, 803–818. 10.1038/s41422-018-0049-z.

29. Molbay, M., Kolabas, Z.I., Todorov, M.I., Ohn, T.L., and Erturk, A. (2021). A guidebook for DISCO tissue clearing. Mol Syst Biol 17, e9807. 10.15252/msb.20209807.

30. Gao, R., Asano, S.M., Upadhyayula, S., Pisarev, I., Milkie, D.E., Liu, T.L., Singh, V., Graves, A., Huynh, G.H., Zhao, Y., et al. (2019). Cortical column and whole-brain imaging with molecular contrast and nanoscale resolution. Science 363. 10.1126/science.aau8302.

31. Tavakoli, M.R., Lyudchik, J., Januszewski, M., Vistunou, V., Agudelo Duenas, N., Vorlaufer, J., Sommer, C., Kreuzinger, C., Oliveira, B., Cenameri, A., et al. (2025). Light-microscopy-based connectomic reconstruction of mammalian brain tissue. Nature 642, 398–410. 10.1038/s41586-025-08985-1.

32. Erturk, A. (2024). Deep 3D histology powered by tissue clearing, omics and AI. Nat Methods 21, 1153–1165. 10.1038/s41592-024-02327-1.

33. Kolabas, Z.I., Kuemmerle, L.B., Perneczky, R., Forstera, B., Ulukaya, S., Ali, M., Kapoor, S., Bartos, L.M., Buttner, M., Caliskan, O.S., et al. (2023). Distinct molecular profiles of skull bone marrow in health and neurological disorders. Cell 186, 3706–3725 e3729. 10.1016/j.cell.2023.07.009.

34. Chu, N.T.L., Dregval, O., Chang, Y.W., Kriukov, E., Tian, X., Liu, X., Trompet, D., Zhang, M.S., Li, L., Li, Z., et al. (2026). Three-dimensional quantitative tissue clearing reveals differences in osteovascular niche of aged and young human mesenchymal stromal cells. Nat Biomed Eng. 10.1038/s41551-026-01645-3.

35. Voigt, F.F., Kirschenbaum, D., Platonova, E., Pages, S., Campbell, R.A.A., Kastli, R., Schaettin, M., Egolf, L., van der Bourg, A., Bethge, P., et al. (2019). The mesoSPIM initiative: open-source light-sheet microscopes for imaging cleared tissue. Nat Methods 16, 1105–1108. 10.1038/s41592-019-0554-0.

36. Konold, P.E., Yoon, E., Lee, J., Allen, S.L., Chapagain, P.P., Gerstman, B.S., Regmi, C.K., Piatkevich, K.D., Verkhusha, V.V., Joo, T., and Jimenez, R. (2016). Fluorescence from Multiple Chromophore Hydrogen-Bonding States in the Far-Red Protein TagRFP675. J Phys Chem Lett 7, 3046–3051. 10.1021/acs.jpclett.6b01172.

37. Coppola, F., Perrella, F., Petrone, A., Donati, G., and Rega, N. (2020). A Not Obvious Correlation Between the Structure of Green Fluorescent Protein Chromophore Pocket and Hydrogen Bond Dynamics: A Choreography From ab initio Molecular Dynamics. Front Mol Biosci 7, 569990. 10.3389/fmolb.2020.569990.

38. Foroughipour, S.M., Becker, K., Foroughipour, M., Ghaffari Tabrizi-Wizsy, N., Sarem, N., Fuchssteiner, C., and Saghafi, S. (2024). Converting a symmetrical Gaussian beam into a thin tunable light sheet. Methods in Microscopy 1, 65–75. 10.1515/mim-2024-0006

39. Vladimirov, N., Voigt, F.F., Naert, T., Araujo, G.R., Cai, R., Reuss, A.M., Zhao, S., Schmid, P., Hildebrand, S., Schaettin, M., et al. (2024). Benchtop mesoSPIM: a next-generation open-source light-sheet microscope for cleared samples. Nat Commun 15, 2679. 10.1038/s41467-024-46770-2.

40. Cregg, J.M., Leiras, R., Montalant, A., Wanken, P., Wickersham, I.R., and Kiehn, O. (2020). Brainstem neurons that command mammalian locomotor asymmetries. Nat Neurosci 23, 730–740. 10.1038/s41593-020-0633-7.

41. Usseglio, G., Gatier, E., Heuze, A., Herent, C., and Bouvier, J. (2020). Control of Orienting Movements and Locomotion by Projection-Defined Subsets of Brainstem V2a Neurons. Curr Biol 30, 4665–4681 e4666. 10.1016/j.cub.2020.09.014.

42. Truckenbrodt, S., Sommer, C., Rizzoli, S.O., and Danzl, J.G. (2019). A practical guide to optimization in X10 expansion microscopy. Nat Protoc 14, 832–863. 10.1038/s41596-018-0117-3.

43. Mai, H., Wang, Y., Zhu, Y., Zhao, Y., Chen, Y., Hoeher, L., Al-Maskari, R., Luo, J., and Erturk, A. (2026). Whole-mouse immunolabeling at cellular resolution for comprehensive 3D atlases. Nat Protoc. 10.1038/s41596-026-01363-9.

44. Tainaka, K., Kubota, S.I., Suyama, T.Q., Susaki, E.A., Perrin, D., Ukai-Tadenuma, M., Ukai, H., and Ueda, H.R. (2014). Whole-body imaging with single-cell resolution by tissue decolorization. Cell 159, 911–924. 10.1016/j.cell.2014.10.034.

45. Stefaniuk, M., Gualda, E.J., Pawlowska, M., Legutko, D., Matryba, P., Koza, P., Konopka, W., Owczarek, D., Wawrzyniak, M., Loza-Alvarez, P., and Kaczmarek, L. (2016). Light-sheet microscopy imaging of a whole cleared rat brain with Thy1-GFP transgene. Sci Rep 6, 28209. 10.1038/srep28209.

46. Gomez-Gaviro, M.V., Sanderson, D., Ripoll, J., and Desco, M. (2020). Biomedical Applications of Tissue Clearing and Three-Dimensional Imaging in Health and Disease. iScience 23, 101432. 10.1016/j.isci.2020.101432.

47. Henning, Y., Osadnik, C., and Malkemper, E.P. (2019). EyeCi: Optical clearing and imaging of immunolabeled mouse eyes using light-sheet fluorescence microscopy. Exp Eye Res 180, 137–145. 10.1016/j.exer.2018.12.001.

48. Klingberg, A., Hasenberg, A., Ludwig-Portugall, I., Medyukhina, A., Mann, L., Brenzel, A., Engel, D.R., Figge, M.T., Kurts, C., and Gunzer, M. (2017). Fully Automated Evaluation of Total Glomerular Number and Capillary Tuft Size in Nephritic Kidneys Using Lightsheet Microscopy. J Am Soc Nephrol 28, 452–459. 10.1681/ASN.2016020232.

49. Capelli, P., Pivetta, C., Soledad Esposito, M., and Arber, S. (2017). Locomotor speed control circuits in the caudal brainstem. Nature 551, 373–377. 10.1038/nature24064.

50. De Zeeuw, C.I., Simpson, J.I., Hoogenraad, C.C., Galjart, N., Koekkoek, S.K., and Ruigrok, T.J. (1998). Microcircuitry and function of the inferior olive. Trends Neurosci 21, 391–400. 10.1016/s0166-2236(98)01310-1.

51. Na, M., Kim, K., Oh, K., Choi, H.J., Ha, C., and Chang, S. (2022). Sodium Cholate-Based Active Delipidation for Rapid and Efficient Clearing and Immunostaining of Deep Biological Samples. Small Methods 6, e2100943. 10.1002/smtd.202100943.

52. Kim, K., Lee, K., Kang, T., Lee, J., Lee, W., Lee, J.Y., and Chang, S. (2025). A novel protein-preserving passive tissue clearing approach using sodium cholate and urea for whole-organ imaging. Exp Mol Med 57, 2292–2304. 10.1038/s12276-025-01550-w.

53. Murakami, T.C., Xia, M., Maeda, Y., Yin, Y., Barbano, P.E., Lin, Z., Mano, T., Tainaka, K., Reiter, S., and Heintz, N. (2026). Artificial intelligence-driven whole-brain cell mapping with highly multiplexed in situ hybridization. Neuron 114, 1380–1398 e1389. 10.1016/j.neuron.2025.12.027.

54. Challis, R.C., Ravindra Kumar, S., Chan, K.Y., Challis, C., Beadle, K., Jang, M.J., Kim, H.M., Rajendran, P.S., Tompkins, J.D., Shivkumar, K., et al. (2019). Systemic AAV vectors for widespread and targeted gene delivery in rodents. Nat Protoc 14, 379–414. 10.1038/s41596-018-0097-3.

55. Alpar, A., Zahola, P., Hanics, J., Hevesi, Z., Korchynska, S., Benevento, M., Pifl, C., Zachar, G., Perugini, J., Severi, I., et al. (2018). Hypothalamic CNTF volume transmission shapes cortical noradrenergic excitability upon acute stress. EMBO J 37. 10.15252/embj.2018100087.

56. Korchynska, S., Rebernik, P., Pende, M., Boi, L., Alpar, A., Tasan, R., Becker, K., Balueva, K., Saghafi, S., Wulff, P., et al. (2022). A hypothalamic dopamine locus for psychostimulant-induced hyperlocomotion in mice. Nat Commun 13, 5944. 10.1038/s41467-022-33584-3.

57. Paxinos, G., and Franklin, K.B.J. (2019). Paxinos and Franklin’s The mouse brain in stereotaxic coordinates, 5th Edition (Academic Press, Elsevier).

58. Saghafi, S., Haghi-Danaloo, N., Becker, K., Sabdyusheva, I., Foroughipour, M., Hahn, C., Pende, M., Wanis, M., Bergmann, M., Stift, J., et al. (2018). Reshaping a multimode laser beam into a constructed Gaussian beam for generating a thin light sheet. J Biophotonics 11, e201700213. 10.1002/jbio.201700213.

59. Mai, H., Luo, J., Hoeher, L., Al-Maskari, R., Horvath, I., Chen, Y., Kofler, F., Piraud, M., Paetzold, J.C., Modamio, J., et al. (2024). Whole-body cellular mapping in mouse using standard IgG antibodies. Nat Biotechnol 42, 617–627. 10.1038/s41587-023-01846-0.

60. Blain, R., Couly, G., Shotar, E., Blevinal, J., Toupin, M., Favre, A., Abjaghou, A., Inoue, M., Hernandez-Garzon, E., Clarencon, F., et al. (2023). A tridimensional atlas of the developing human head. Cell 186, 5910–5924 e5917. 10.1016/j.cell.2023.11.013.

